# Integrin blocking peptide reverses immunosuppression in experimental gliomas and improves anti-PD-1 therapy outcome

**DOI:** 10.1101/2024.08.06.606798

**Authors:** Aleksandra Ellert-Miklaszewska, Paulina Pilanc-Kudlek, Katarzyna Poleszak, Adria-Jaume Roura, Salwador Cyranowski, Mitrajit Ghosh, Szymon Baluszek, Maria Pasierbinska, Bartłomiej Gielniewski, Julian Swatler, Yuliana Hovorova, Kamil Wojnicki, Bozena Kaminska

**Author notes:** Corresponding authors: Ellert-Miklaszewska A., Kaminska B. These authors contributed equally: Ellert-Miklaszewska A., Pilanc-Kudlek P.

## Abstract

Immune checkpoint inhibitors (ICI) presented clinical benefits in many cancer patients but invariably fail in glioblastoma (GBM), the most common and deadly primary brain tumor. Lack of ICI efficacy in GBM is attributed to the accumulation of immunosuppressive myeloid cells that create the “cold” tumor microenvironment (TME) impeding infiltration and activation of effector T cells. We developed a designer RGD peptide that hindered glioma-instigated, integrin-mediated pro-tumoral reprogramming of myeloid cells and blocked microglia-dependent invasion of human and mouse glioma cells in co-cultures *in vitro*. Intratumorally-delivered RGD alone did not reduce glioma growth in syngeneic mice but prevented the emergence of immunosuppressive myeloid cells and led to peritumoral blood vessels normalization. Furthermore, combining RGD with immunotherapy using PD-1 blockade reduced tumor growth, led to upsurge of proliferating, interferon-ɣ producing CD8^+^T cells and depleted regulatory T cells. Transcriptomic profiles of myeloid cells were altered by the combined treatment, consistently with the restored “hot” inflammatory TME and boosted immunotherapy responses. RGD modified the phenotypes of myeloid cells in human gliomas in nude mice. Thus, combining the integrin blockade with ICI reinvigorates antitumor immunity and paves the way to improve immunotherapy outcomes in GBM.

## INTRODUCTION

Immune checkpoint inhibitors (ICIs) have altered the therapy outcomes in a wide range of cancers, providing unparalleled survival benefits in some patients. Blocking antibodies against programmed cell death 1 (PD-1, CD279) or programmed cell death-ligand 1 (PD-L1, CD274) (anti-PD-1 or anti-PD-L1, respectively) are employed in more than 20 different indications (1). Unfortunately, only approximately 20% of cancer patients (mostly with melanoma and lung cancer) achieve the long-term, durable responses with ICIs, while resistance is frequent for most patients (2). Glioblastoma (GBM) is the most common and lethal primary brain tumor among adult-type diffuse gliomas (3). Despite some promising results of the phase II/III trials, ICIs have failed in the phase III randomized trials in both newly diagnosed and recurrent GBM patients (4, 5). This limited efficacy of ICIs is attributed to the highly immunosuppressive tumor microenvironment (TME) of GBM with a high contribution of tumor supportive myeloid cells (up to 30% of the tumor mass) (6, 7). Preclinical and clinical data demonstrate numerous immune deficits and tumor immune evasion mechanisms operating in the TME that can impede the efficacy of immunotherapies with ICIs. Experimental studies show that reversing the protumor reprogramming of myeloid cells and normalization of tumor neovasculature may overcome TME-instigated resistance to immunotherapy (8–11).

GBM is known as a “cold” tumor with low mutation burden, abundance of suppressive myeloid cells and poor infiltration of effector T lymphocytes and NK cells (7). Human GBM and murine gliomas attract brain resident microglia and bone marrow-derived myeloid cells, jointly called glioma-associated microglia and macrophages (GAMs) and impose tumor-supportive programs in these cells. GAMs increase tumor cell invasion, angiogenesis and display immunosuppressive functions by inhibiting antitumor T cell proliferation and cytotoxic activity, and initiating signals for their anergy or exhaustion, while promoting the expansion of protumorigenic T regulatory cells (12–15). Defective neoangiogenic blood vessels drive metabolic reprogramming in hypoxic niches and impair trafficking of T cells to the tumor, which aggravates functional deficits in antitumor immunity and eventually contributes to ICIs therapy failure (16–18). Multiple strategies have been proposed to modify glioblastoma TME for better therapeutic responses, including targeting GAMs with an inhibitor of the colony-stimulating factor (CSF)-1 receptor (19) or CCR2/CCL2 axis blockade (9). Strategies that render the TME “hot” may sensitize these tumors to concurrent or subsequent immunotherapy.

We have previously identified glioma derived osteopontin/secreted phosphoprotein 1 (SPP1) and lactadherin/milk fat globule-epidermal growth factor 8 (MFG-E8) as activators of myeloid cell reprogramming in gliomas (20). SPP1 and MFG-E8 bind to the αvβ3 and αvβ5 integrin receptors on myeloid cells via the RGD motif (arginine–glycine–aspartate). MFG-E8 promotes vascular endothelial growth factor (VEGF)-mediated neoangiogenesis via αvβ3/αvβ5 that are overexpressed on newly formed tumor vessels in murine and human gliomas (21, 22). Congruently, SPP1 via αvβ5 recruited GAMs to mouse gliomas and systemic administration of osteopontin-specific aptamers increased median animal survival time by 68% (23). Silencing of *Spp1* or *Mfge8* expression in glioma cells reduced tumor growth and prevented GAMs reprogramming in orthotopic rat gliomas (20). Similarly, knockdown of GBM-secreted periostin (POSTN), another integrin ligand driving recruitment of GAMs, had anti-glioma effects (24). We designed a synthetic 7-amino-acid peptide inclosing RGD motif (‘RGD’), which competitively blocks the interactions of SPP1 and MFG-E8 with integrins, and prevents the reprogramming of microglia induced by glioma secretome. Pretreatment with the RGD peptide abolished the activation of FAK-dependent signaling pathway, microglial phagocytosis and migration, and reduced the expression of proinvasive phenotype genes in microglia (20).

Herein, we demonstrate that RGD prevents microglia activation and reduces microglia-dependent invasion of murine and human glioma cells *in vitro*. Intratumorally delivered RGD as a monotherapy effectively blocks the reprogramming of GAMs and shifts the glioma TME towards a “hot” phenotype but does not reduce tumor growth. Application of the RGD peptide alters immune cell composition and cell functionalities augmenting proinflammatory responses, normalizes tumor vasculature, and reduces tumor growth when co-administered with anti-PD-1 immunotherapy. We provide the evidence that combining the integrin blockade with ICI reinvigorates antitumor immunity by boosting both innate and adaptive immune responses, and indicates a novel strategy to enhance the efficacy of immunotherapy. These results open a new perspective for treatment of GBM patients and pave the way to improve responses to ICIs in other cancers.

## RESULTS

### RGD peptide blocks microglia-induced invasion of murine and human glioma cells

The integrin blocking peptide (RGD) was designed to block the binding of glioma-derived SPP1 and MFGE8 to αvβ3/αvβ5-integrins (*20*). Interactome analysis of the human integrin αv generated by STRING (Fig. 1A) depicted SPP1 and MFG-E8 among top 10 proteins with experimentally confirmed physical associations with these integrins, via complexes formed with β3 and β5 subunits. Analyzing publicly accessible scRNA-seq data from human gliomas (25) via the SingleCell portal we found the predominant expression of *ITGAV*, *ITGB3* and *ITGB5* (αv, β3 and β5 integrin encoding genes, respectively) in clusters of myeloid cells, pericytes and endothelial cells (Fig. 1B). This indicates that glioma-associated myeloid cells and blood vessels might be the main targets of integrin blockade with the RGD peptide. Moreover, to validate the potential of RGD based therapy, we explored TCGA expression datasets of low- and high-grade gliomas. The levels of *ITGAV*, *ITGB3* and *ITGB5* were significantly higher in GBM than in non-tumor samples (Fig. S1A). *ITGAV* and *ITGB5* expression was also elevated in astrocytomas. Among all GBM, the expression of these genes was the highest in the mesenchymal GBM subtype associated with the worst clinical outcome (Fig. S1B). Survival analysis based on the expression of individual integrin genes (Fig. S1C) or a multi-gene signature (Fig. 1C) revealed that high levels of *ITGAV*, *ITGB3* and *ITGB5* correlate with a poor prognosis in glioma patients. This data provided the rationale for further exploration of integrin blocking peptide efficacy in the treatment of GBM.

**Fig. 1.**
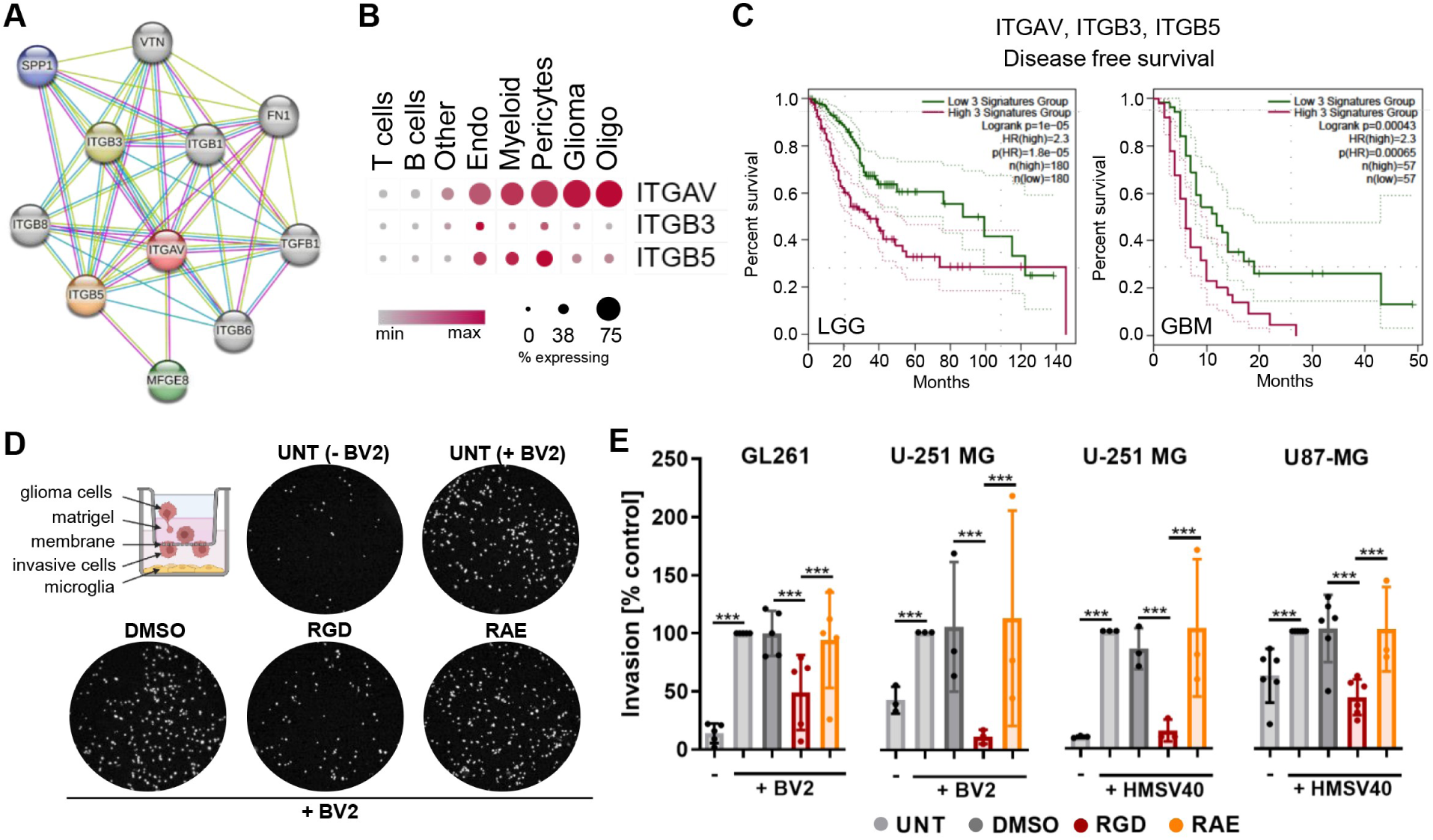
The RGD peptide abrogates microglia-dependent glioma cell invasion. (A) Top 10 predicted associations of integrin αv (ITGAV)-interacting proteins modeled using STRING. Purple line - experimental evidence, yellow line – text mining evidence, light blue line - database evidence. (B) Scaled mean expression of integrin-coding genes in indicated cell types in scRNA-seq of 44 human gliomas (study GSE182109, SingleCell portal). (C) Kaplan-Meier curves generated using GEPIA2 for low-grade glioma (LGG) and GBM patients with high (in purple) and low (in green) expression status of a multi-gene signature (cutoff-high 65% and cutoff-low 35%; dotted line depicts 95% CI. (D-E) Matrigel invasion assay. Glioma cell invasion in the absence or presence of microglial cells (see the scheme) was measured after 18 h of co-culture. (D) Representative images of DAPI-stained nuclei of invading GL261 cells in co-culture with BV2 microglia; cells were untreated (UNT) and treated with solvent (DMSO, 0.2%), 100 µM RGD or RAE peptides. (E) Quantification of GL261, U-251 MG and U87-MG glioma cells invasion co-cultured with murine BV2 or human HMSV40 microglial cells, in the presence of DMSO, RGD or RAE peptides. Invasion of untreated glioma cells (UNT) co-cultured with microglia cells was set as 100%. All experiments were performed three times, in duplicates. Statistical significance was calculated using the chi-square test;***p value 0.001.

Glioma-activated microglia stimulate glioma invasion *in vitro*, in organotypic brain slices and *in vivo* (20, 26–28). We evaluated the effect of the designed RGD peptide on microglia-dependent invasion of murine (GL261) and human (U251-MG, U87-MG) glioma cells. Co-culture of glioma cells with mouse BV2 or human HMSV40 microglial cells, enhanced invasion of glioma cells through the Matrigel layer (Fig. 1D-E). Representative images of invading GL261 cells in the absence or presence of BV2 microglia in the control conditions and with RGD or a control RAE peptide are shown in Fig. 1D. In the RAE peptide, the RGD motif (arginine-glycine-aspartic acid) is changed to RAE (arginine-alanine-glutamic acid), which blunts its activity. Only the RGD peptide (and not RAE) significantly reduced the microglia-induced invasion of glioma cells and was effective in all cell culture settings (including human glioma cells/murine- or human microglia co-cultures) (Fig. 1E).

The peptides exhibited no cytotoxic effects on glioma or microglial cells, as evidenced by MTT metabolism and BrdU incorporation assays. Viability of cells cultured with RGD was only slightly reduced in comparison to untreated or solvent-treated cells (Fig. S2A). Cell proliferation was not affected by RGD in cell cultures, except for U87-MG cells, in which the average proliferation was insignificantly lower compared to controls (Fig. S2B). Adding the RAE peptide had no effects on cell viability and proliferation (Fig. 2A-B). These data confirm that the inhibitory effect of the RGD peptide on microglia-supported glioma invasion is due to the peptide interference in glioma-microglia communication and reprogramming.

**Fig. 2.**
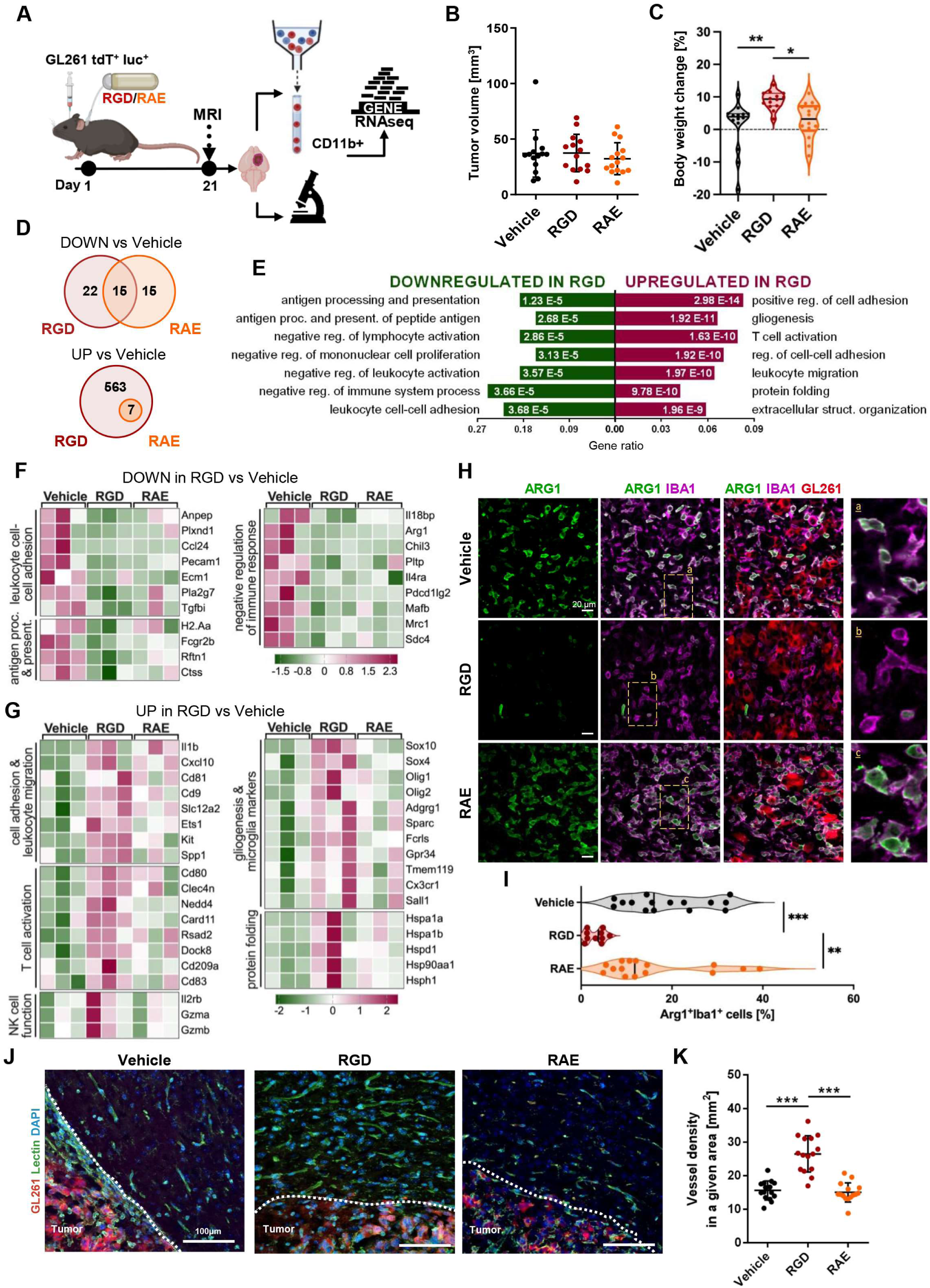
Intratumorally-delivered RGD modifies tumor microenvironment. (A) Schematic representation of the experimental pipeline. (B) Mice implanted with GL261 tdTomato+luc+ glioma cells received water (Vehicle), RAE or RGD peptide via osmotic pumps. Tumor volume was quantified by MRI at 21 DPI, n=15-16 per group. (C) Comparison of mice body weight between 0 and 21 DPI; n=15-16; the line is plotted at the mean. (D-G) RNA-seq gene expression profiling of CD11b^+^ cells from the tumor-bearing hemispheres of mice at 21 DPI. (D) Venn diagrams showing a number of down - and upregulated DEGs in CD11b^+^ cells from RGD and RAE compared to the vehicle group. (E) Functional enrichment analysis with Gene Ontology (GO) biological processes for up- and downregulated genes in RGD compared to Vehicle. Enriched GO pathways (selected from top ranked) are shown, the size of the bars indicates gene ratios (a number of genes annotated to the pathway/total number of DEGs with adjusted p-values < 0.05). Z-score heatmaps for selected down- (F) and upregulated (G) genes in the CD11b^+^ cells from RGD vs Vehicle treated animals. (H) Representative images of the glioma-bearing brains (at 21 DPI) stained with anti-Iba1 (in purple) and anti-Arg1 (in green) antibodies, and co-stained with DAPI are shown. (I) Quantification of IBA1^+^Arg1^+^ cells. Average values from 3 fields are presented; n=5. (J) Representative images of the sections from glioma bearing brains stained with lectin (green) and co-stained with DAPI are shown. (K) Quantification of vessel density. Three independent sections from each mouse brain were analyzed; in total 15 sections in each group. Each dot represents an average density in 2 regions of interest (ROI, see Fig. S4D), both near the tumor core. Significance was calculated with One-Way ANOVA, followed by Tukey’s post-hock test.*p < 0.05; **p < 0.01; ***p < 0.001; ****p < 0.0001. Data in dotplots are presented as mean ± SD. Line on the violin plots depicts the mean.

### RGD alleviates the emergence of the tumor-supportive TME in gliomas

Next, we sought to determine whether the RGD peptide affects the reprogramming of myeloid cells by gliomas *in vivo* and if it interferes with tumor growth in mice. GL261 cells implanted into the striatum of immunocompetent C57BL/6J mice recapitulate many characteristics of human GBMs, and are frequently used in the preclinical glioma research (29). The peptide was administered intratumorally via osmotic micropumps, installed at the time of glioma cells implantation (Fig. S3A).

Mass spectrometry analysis of the RGD peptide incubated in the osmotic pumps at 37°C did not reveal any products of degradation or any decline in the peptide level throughout the incubation period, showing stability of RGD in water solution and no binding to the pump itself (Fig. S3B). According to the biodistribution studies, an intratumorally administered FITC-labeled RGD peptide was present in the tumor core and diffused to the surrounding tissue, indicating efficient delivery and penetration within the brain parenchyma (Fig. S3C).

For the evaluation of antitumor activity, RGD, RAE or vehicle (water) were delivered to GL261 tdTomato^+^Luc^+^ gliomas via osmotic pumps for 21 days (Fig. 2A). According to magnetic resonance (MRI) scans at 21 day post-implantation (DPI), the tumor growth was not affected by the peptides as compared to controls (Fig. 2B). However, the weight of RGD-treated mice significantly increased (Fig. 2C), which is indicative of animal well-being in experimental tumor models. Meanwhile, the animals which received the vehicle or RAE lost their weight over the time or their weight remained unchanged.

To determine the effects of the RGD peptide on tumor infiltrating myeloid cells *in vivo*, we compared the transcriptomic profiles of FACS-sorted glioma-associated CD11b^+^ cells from all experimental groups. At 21 DPI, percentages of intratumoral CD45^low^CD11b^+^ microglia and CD45^high^CD11b^+^ monocytes/macrophages were similar across treatments (Fig. S4A) However, there were numerous transcriptomic differences reflecting changes in the functionalities of the cells. Among differentially expressed genes (DEGs), downregulated DEGs showed some overlap between both peptide treatments, however out of 563 upregulated genes in the RGD group, only 7 were identified as DEGs in CD11b^+^ cells from RAE-treated mice, which confirms the specificity of response to the RGD peptide (Fig. 2D).

Gene Ontology (GO: Biological Process) enrichment analysis (Fig. 2E) indicated that the downregulated genes in the RGD group were related to antigen processing and presentation, negative regulation of lymphocyte activation and negative regulation of immune system processes. Among the differentially upregulated genes in the RGD group, we found a significant overrepresentation of genes involved in regulation of cell adhesion, T cell activation and leukocyte migration.

The downregulated genes in the RGD group (Fig. 2F), included those involved in leukocyte cell-cell adhesion such as *Anpep*, which encodes a membrane bound enzyme promoting monocyte adhesion to the endothelium during leukocyte exit into the tissues (30), and *Pecam1,* a member of the immunoglobulin superfamily, engaged in leukocyte migration, angiogenesis, and integrin activation. Moreover, some of these genes encode factors involved in the macrophage-driven suppression of immune responses such as *Arg1*, *Chil3* and *Mrc1*. Additionally, decreased levels of *Pdcd1lg2*, *Il4ra* and *Il18bp* were observed. Binding of programmed cell death 1 ligand 2 (PD-L2, encoded by *Pdcd1lg2)* to its receptor PD-1 leads to inhibition of T cell proliferation and inflammatory cytokine production. *Il4ra* encodes the alpha chain of the receptor for interleukin (IL)-4, which is well known for skewing macrophages toward the alternatively-activated phenotype. IL-18 binding protein (coded by *Il18bp*) is an inhibitor of the proinflammatory cytokine IL18. Treatment with the RGD peptide resulted in reduced expression of genes coding for proteins related to antigen processing and presentation, including *H2-Aa* encoding major histocompatibility complex class II (MHC-II) protein, and *Ctss* coding for Cathepsin S which participates in the degradation of antigenic proteins to peptides for presentation on MHC-II molecules.

CD11b^+^ from the RGD treated mice showed increased expression of genes related to T cell activation (Fig. 2G). This group included *CD80,* encoding co-stimulatory molecule necessary for T cell activation, and *Rsad2,* which is an interferon-stimulated gene involved in IRF7-mediated maturation of mouse dendritic cell (DC) and their antitumor efficacy. In parallel, genes coding for dedicator of cytokinesis 8 (*Dock8*) and DC-specific C-type lectin receptor DC-SIGN (*CD209a*) were upregulated; these proteins may augment the adaptive immune response by promoting activation of T cells. Moreover, in CD11b^+^ cells from mice treated with RGD, genes coding for: NEDD4 – E3 ligase involved in reprogramming of tumor associated macrophages via CSF1R degradation (31), CD81 - a regulator of the recruitment of NK cells (32), CD9 - a member of the transmembrane 4 superfamily (TM4SF) involved in cell growth, adhesion and motility, cytokine Il1b and Cxcl10 - IFNγ-stimulated chemokine that attracts T cells through binding to its cognate receptor CXCR3, showed increased expression. Genes associated with cytotoxic activity of NK cells were significantly upregulated after the RGD treatment, including genes coding for interleukin 2 receptor subunit beta (*IL2rb*) involved in T cell immune responses, as well as granzyme A (*Gzma*) and granzyme B (*Gzmb*), which are the major cytolytic factors in antitumor response of cytotoxic NK cells. In addition, RGD triggered the overexpression of *Fcrls* and *Tmem119,* well known homeostatic microglia markers, and *Gpr34*, which is upregulated in microglia during inflammation. Another important group of upregulated genes in CD11b^+^ cells from RGD-treated mice included those coding for several heat shock proteins, which have been implicated in the stimulation of both innate and adaptive immunity (33).

To track cell type alterations induced within the brain CD11b^+^ cells by peptide treatments, we deconvoluted bulk RNA-seq gene expression data leveraging CITE-seq transcriptomic signatures of subpopulations of CD11b^+^ cells from naïve and glioma-bearing brains (34) (Fig. S4B-C). The predicted frequencies of homeostatic microglia increased in response to the RGD treatment as compared to the vehicle-treated group and those of activated microglia were decreased, although the latter change was at the boundaries of statistical significance. The switch in proportions of homeostatic (the most prevalent in naïve mice) and activated microglia upon RGD indicated the efficacy of the peptide in mitigating tumor-induced reprogramming.

To validate the observed transcriptomic changes, suggestive of a switch of GAMs phenotype upon the RGD administration, we performed immunohistochemical co-staining for Iba1 (a marker of myeloid cells) and Arg1 (a hallmark marker of immunosuppressive/proinvasive GAMs) on the brain sections from glioma-bearing mice. RGD significantly reduced the accumulation of Iba1^+^Arg1^+^ positive cells in the tumors in comparison to the vehicle or RAE groups (Fig. 2H-I). This data supports the findings obtained from the RNA-seq analysis. Although RGD alone did not reduce the tumor volume nor affected the abundance of myeloid cells, it blocked reprogramming of CD11b^+^ cells into proinvasive, immunosuppressive GAMs.

Another feature strongly associated with the TME is the abnormal neovasculature. The structure of blood vessels is disrupted in tumors leading to the formation of numerous small vessels, which are leaky, collapsed, and disorganized, leading to impaired T cell trafficking, appearance of hypoxic niches and poor drug delivery (16). Pathological angiogenesis in GBM is associated with upregulation of the expression of certain integrins, including αvβ3 and αvβ5. For these reasons, we evaluated the density of vasculature in GL261 glioma-bearing mice receiving intratumorally RGD or RAE peptides for 21 days. Lectin staining revealed numerous microvessels in the peritumoral area of vehicle- and RAE-treated tumor-bearing brains (Fig. 2J-K, Fig. S4D). In the RGD group, the staining and its quantification demonstrated an increased density of large, elongated vessels in the peritumoral area consistent with their improved integrity. These findings suggest that the RGD treatment results in vessel normalization. Overall, the phenotype alterations observed in myeloid cells and improved vasculature organization are indicative of a transition from a “cold” to “hot” TME, primed for complete restoration of antitumor immunity with an adjuvant treatment.

### Combining RGD with PD-1 blockade reduces glioma growth in mice due to augmented antitumor immune responses

Limited efficacy of immune checkpoint inhibitors in GBM is mainly attributed to the highly immunosuppressive TME and altered tumor vasculature impairing T cell trafficking (16). RGD treatment blocked the intratumoral reprogramming of myeloid cells into proinvasive and immunosuppressive GAMs and led to blood vessels normalization. These results prompted us to study if RGD-induced TME and vasculature changes could improve immunotherapy with the anti-PD-1 antibody. The intratumorally delivered RGD peptide was combined with 4 *i.p.* injections of anti-PD-1 (aPD-1, or an isotype control IgG antibody) in immunocompetent mice implanted with GL261 cells (Fig. 3A). MRI scans revealed significantly reduced tumor volumes at 28 DPI (vehicle vs RGD+aPD-1: *p* = 0.0128, r = 0.67) (Fig. 3B-C). In concordance with the observed reduction of tumor burden, the body weight of 75% of mice exposed to the combination treatment for 28 days increased over time, while in the RGD or aPD-1 groups the range of changes of animal weights was similar as in the vehicle group (Fig. 3D).

**Fig. 3.**
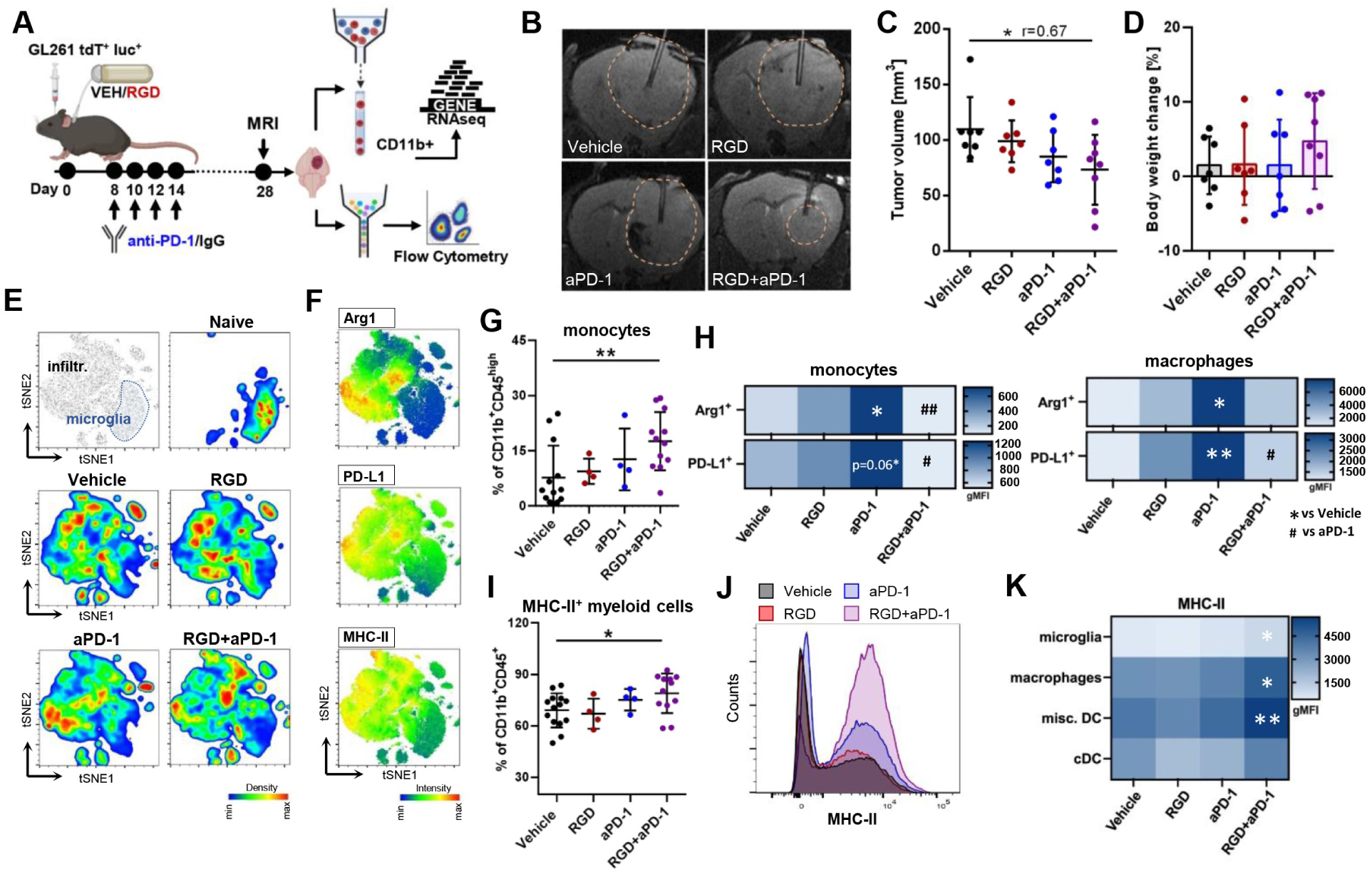
Combining RGD and anti-PD-1 reduces GL261 glioma growth and instigates changes in the myeloid compartment. (A) Scheme of experiments combining RGD delivery with PD-1 blockade. Mice implanted with GL261 tdTomato+luc+ glioma cells received water (vehicle) or the RGD peptide via osmotic pumps for 28 days and anti-PD-1 antibody (aPD-1, 10 mg/kg) or control IgG at day 8, 10, 12 and 14 by i.p. injection. (B-C) Tumor volumes were measured by MRI at 28 DPI. The effect size of treatment was assessed with factorial ANOVA; Vehicle-RGD+aPD-1= 0.67; followed by post-hoc Sidak multiple comparisons test: *p*(Vehicle-RGD+aPD-1)=0.0128, n=7-8. (D) Comparison of body weight in mice between day 0 and 28 DPI; n=7-8; whiskers represent the min-max values, the box extends from the 25th to 75th percentiles, the line is plotted at the median. (E) Unsupervised t-SNE clustering of CD45^+^CD11b^+^ from brains of naïve and Vehicle, RGD, aPD-1 and RGD+aPD-1 treated mice. Microglia and infiltrating myeloid cells are separated by manual gating (the first plot from the left). Pseudocolor smooth density plots depict cell clusters of high and low density in red and blue, respectively. (F) Heatmap plots show the levels of functional markers in a blue-to-red scale (low-high). (G) Percentages of monocytes (CD45^high^CD11b^+^Ly6C^high^F4/80^low^) in tumor-bearing brains. (H) Levels (gMFI) of Arg1 and PD-L1 in glioma-associated monocytes (CD45^high^CD11b^+^Ly6C^high^F4/80^low^) and macrophages (CD45^high^CD11b^+^Ly6C^low^F4/80^high^); n=4-14. (I) Percentages of MHC-II-expressing myeloid cells in tumor-bearing brains and (J) a representative histogram. (K) Levels (gMFI) of MHC-II in glioma-associated monocytes (CD45^high^CD11b^+^Ly6C^high^F4/80^low^), macrophages (CD45^high^CD11b^+^Ly6C^low^F4/80^high^), classical dendritic cells (cDC; CD45^high^CD11b^-^CD11c^+^) and other DC subsets (misc. DC; CD45^high^CD11b^+^CD11c^+^). Data in all quantitative panels are presented as mean ± SD *p < 0.05; **p < 0.01; ***p < 0.001; Statistical analysis by Mann-Whitney test or one-way ANOVA, followed by Tukey multiple comparison test.

Multiparameter flow cytometry analysis was performed on tumor-infiltrating myeloid and lymphoid immune cell populations to tackle changes in the TME composition. At 28 DPI, the total frequency of myeloid cells among all CD45^+^ tumor-infiltrating immune cells as well as the percentages of microglia (CD11b^+^CD45^low^) and infiltrating peripheral myeloid cells (CD11b^+^CD45^high^) did not differ between the vehicle and other experimental groups (Fig. S5A-B). However, t-SNE visualization showed the phenotypic shifts in the myeloid compartment upon different treatments (Fig. 3E-F, Fig. S5C). We noticed a significant increase of monocytes (Ly6C^high^F4/80^low^) within the CD11b^+^CD45^high^ population upon RGD+aPD-1 treatment (Fig. 3G). Anti-PD-1 administration led to increased Arg1 and PD-L1 levels in myeloid cells, while the combination therapy downregulated Arg1 and PD-L1 levels both in monocytes (Ly6C^high^F4/80^low^) and macrophages (Ly6C^low^F4/80^high^) (Fig. 3H). In addition, we observed an increased frequency of MHC-II^+^ cells within the myeloid compartment upon RGD+aPD-1 treatment as compared to the vehicle group (Fig. 3I-J). Upregulated MHC-II levels were detected on microglia, macrophages and on subsets of dendritic cells (DC), mainly CD11b^+^CD11c^+^ DCs but not CD11b^-^CD11c^+^ DCs (Fig. 3K). Augmented MHC-II levels may reflect the improved antigen presenting capacity and facilitate activation of CD8^+^ T cells.

Since detected changes in myeloid cells suggested a proinflammatory switch in the TME of RGD+aPD-1-treated tumors, we investigated how this affected the lymphoid compartment, classically modulated by ICI. The percentage of CD3^+^ T cells among all CD45^+^ tumor-infiltrating immune cells did not differ between the vehicle and other experimental groups (Fig. S5D). However, t-SNE visualizations of flow cytometry data showed quantitative shifts between lymphocyte clusters (Fig. 4A-B, Fig S5E). The frequency of CD8^+^ T cells was increased in anti-PD-1 treated tumors as compared to untreated tumor-bearing mice, and was markedly elevated in the combination group (Fig. 4C). We noticed a significant decrease of the immunosuppressive CD4^+^CD25^high^Foxp3^+^ T regulatory cells (Treg) in the RGD+aPD-1 group. Consequently, the CD8^+^T cells/Treg ratio, which predicts the response to immunotherapy in cancer patients and experimental mouse models, was increased upon RGD+aPD-1 treatment (Fig. 4C). Moreover, ratios of CD8^+^ T cells to CD11b^+^ cells were significantly increased in the RGD+aPD-1 group due to accumulation of cytotoxic T cells, indicating antitumor immune responses in the glioma TME.

**Fig. 4.**
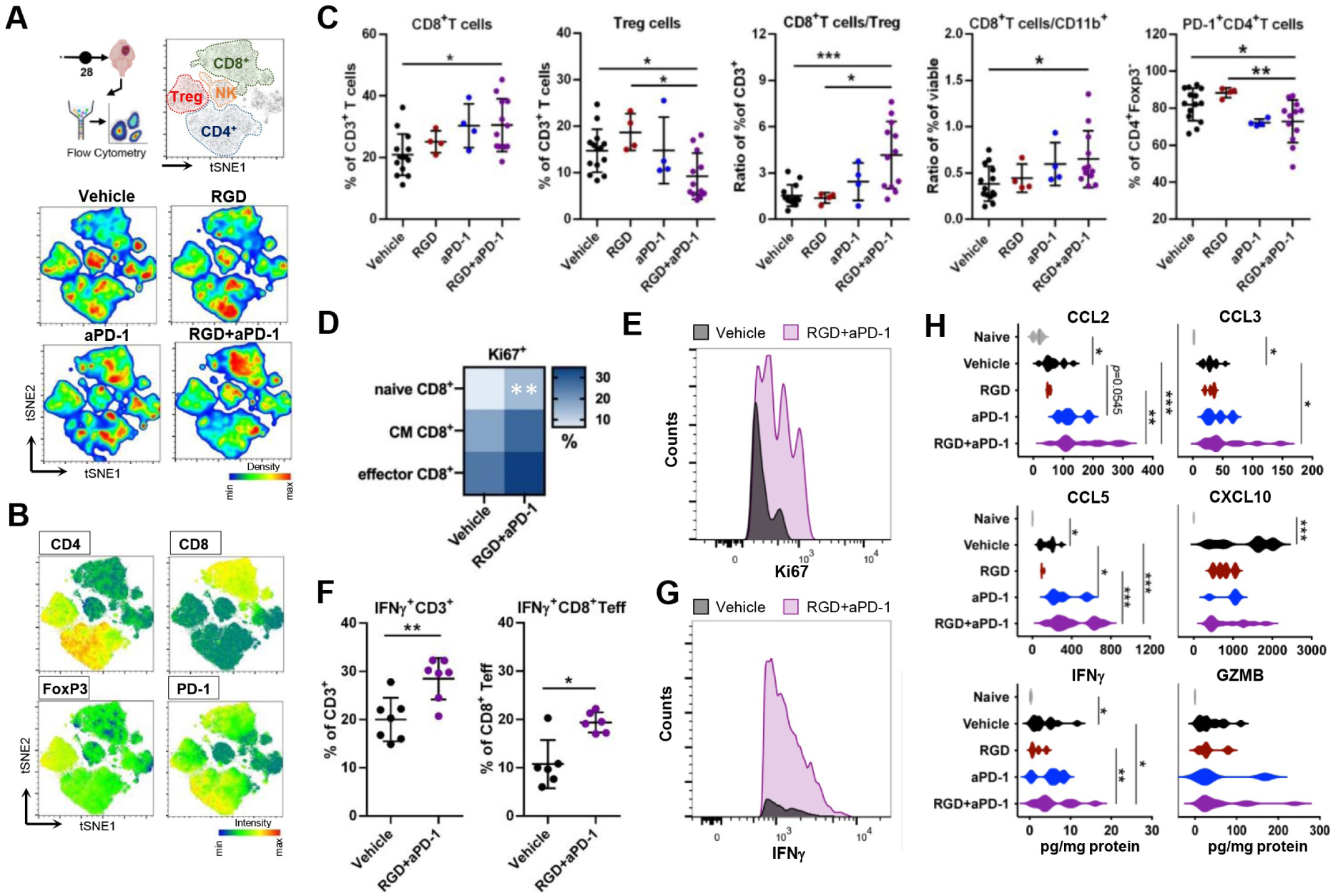
Changes in lymphoid compartments and cytokine levels confer effective antitumor immune responses upon integrin and PD-1 blockade. Experimental workflow - see Fig. 3. (A) Unsupervised t-SNE clustering of CD45^+^CD11b^-^ from the brains of Vehicle, RGD, aPD-1 and RGD+aPD-1 treated mice. The main lymphocyte populations are separated by manual gating (top plot on the right). Pseudocolor smooth density plots depict cell clusters of high and low density in red and blue, respectively. (B) Heatmap plots show the levels of cell identity and functional markers in a blue-to-red scale (low-high). (C) Percentages of CD8^+^ (CD8^+^CD4^-^), Tregs (CD4^+^Foxp3^+^CD25^high^), ratio of CD8^+^/Treg cells, percentage of PD-1^+^CD4^+^Foxp3^-^ T cells and ratio of CD8^+^/CD11b^+^cells isolated from tumor-bearing brains in all experimental groups; n=4-14. (D) Heatmap shows percentages of Ki67^+^ cells among effector (CD44^+^CD62L^-^), central memory (CM; CD44^+^CD62L^+^) and naïve (CD44^-^CD62L^+^) CD8^+^ T cells. (E) A representative histogram of Ki67^+^ effector CD8^+^ T cells. (F) Production of IFNγ by glioma-infiltrating T cells (CD3^+^) and effector CD8^+^ T cells isolated from the brains of Vehicle and RGD+aPD-1 treated mice and stimulated with 50 ng/ml PMA + 1 µg/ml ionomycin in the presence of protein transport inhibitors, n=6. (G) A representative histogram of IFNγ^+^ effector CD8^+^ T cells. Data in all quantitative panels are presented as mean ± SD. Statistical analysis by Mann-Whitney test or one-way ANOVA, followed by Tukey multiple comparison test. (H) The levels of cytokines were determined in brain homogenates of naive and treated mice at 28 DPI using a multiplexed Luminex assay. Violin plots show the levels of tested cytokines in pg/mg of total protein Significance was assessed with One–Way ANOVA and Uncorrected Fisher’s LSD multiple comparison test; N=4-12. ***p < 0.001; **p < 0.01; *p < 0.05.

We also noticed that the percentages of PD-1^+^CD4^+^Foxp3^-^ cells significantly decreased in RGD+aPD-1-treated animals as compared to the peptide-alone or vehicle groups (Fig. 4C). PD-1^+^CD4^+^Foxp3^-^ cells contribute to tumor immune evasion, as they accumulate intratumorally during tumor progression and limit effector CD8^+^ T cell functions. The persistence of elevated frequencies of these cells after PD-1 blockade is a negative prognostic factor in melanoma patients (35).

Tumor-infiltrating T cells usually exhibit a terminally exhausted phenotype, marked by a loss of self-renewal capacity (7). In RGD+aPD-1-treated mice, the average frequencies of Ki67^+^ cells among glioma-infiltrating effector, central memory (CM) and naïve CD8^+^ T cells were increased, indicating augmented proliferation of these cell subsets, with the latter population showing statistically significant increase versus the vehicle controls (Fig. 4D-E).

To study if the effector functions of glioma infiltrating CD8^+^ T cells are restored upon the RGD+aPD-1 treatment, we quantified the production of IFNγ by T cells stimulated *ex vivo* with PMA and ionomycin. CD3^+^ T cells derived from the RGD+aPD-1 tumors, including effector CD8^+^ T cells, showed increased IFNγ production (Fig. 4F-G). Together, the presented findings demonstrate a profound remodeling of the immune landscape of glioma TME after the RGD+aPD-1 treatment with a specific shift in the intratumoral reprogramming of infiltrating CD11b^+^CD45^high^ myeloid populations, which results in the augmented responses of CD8^+^ T cells.

The reinvigoration of antitumor responses at 28 DPI upon RGD+aPD-1 administration was further evaluated by measuring the levels of pro- and anti-inflammatory cytokines in the brain homogenates and sera (Fig. 4H, Fig. S5F-G). C-C Motif Chemokine Ligand (CCL) 2, CCL3 and CCL5 were increased in the brain homogenates from aPD-1 treated mice (vs the vehicle group) and showed even higher levels in RGD+aPD-1 treated animals. CXCL10 showed elevated levels (compared to naïve mice) in the brains of glioma-bearing mice independent of the treatment. IFNɣ, granzyme B and TNFα, which are produced by activated immune cells and linked to antitumor immune responses, were elevated in aPD-1 and RGD+aPD-1 groups as compared to the vehicle, with the levels of IFNɣ, the master immune effector cytokine, being significantly higher in the RGD+aPD-1 group. Interestingly, the levels of these cytokines measured in the sera of the same animals remained unchanged upon the treatments, except from decreased levels of CCL2 in the RGD+aPD-1 group (Fig. S5G). CCL2 and CCL5 are involved in monocyte attraction to tumors (36), CCL5 is crucial for DC recruitment and in conjunction with CXCL9/10 generates main chemotactic ques for effector T cells (37, 38). CCL3 is considered as a marker of proinflammatory macrophages and leads to improved DC and T cell responses in TME (39). The observed contextual changes in the cytokine levels support the notion of restoring the proinflammatory TME in experimental gliomas upon RGD+aPD-1 treatment.

Antitumor immune responses in the aPD-1 and RGD+aPD-1 treated gliomas evolved with time as at 21 DPI the tumor volume was not changed upon the treatment (Fig. S6A-B) and the myeloid compartment was yet dominated by Arg1^+^ and PD-L1^+^ immunosuppressive macrophages (Fig. S6C-D). Interestingly, the observed effects were specific to the tumor niche as the peripheral monocytes and granulocytes, isolated from the spleen, were not affected (Fig. S6E-F). At 21 DPI, there were already some qualitative changes in the lymphoid compartment (Fig. S6G-J). Intratumoral levels of effector CD8^+^ T cells were significantly increased after aPD-1 and RGD+aPD-1 treatments. Concurrently, the percentages of Tregs remained unchanged in the tumor but were elevated at the periphery in aPD-1 and RGD+aPD-1 groups, counteracting the revival of antitumor responses. Similar evolution of responses to ICIs with initial upregulation of immunosuppression as a feedback loop has been previously reported (27, 40).

### GAMs from RGD+anti-PD-1 treated mice are transcriptionally reprogrammed for antitumor responses

Transcriptome profiling of CD11b^+^ cells isolated from tumor-bearing brains of mice at 28 DPI using RNA-seq was performed to determine underlying differences between the groups. The most profound changes versus the vehicle group were detected upon the combination treatment (Fig. 5A). The GO analysis of the DEGs in this group revealed upregulation of several pathways related to inflammatory and antitumor responses, such as T cell activation, cytokine mediated signaling pathway, adaptive immune response, response to interferon gamma, leukocyte migration and antigen processing and presentation (Fig. 5B). These categories comprised genes coding for co-stimulatory molecules and markers delineating mature APC functions (*Icam1, CD40, CD86*), chemokines, cytokines and their receptors (*Ccl7, Ccl5, Ccl2, Cxcl9, Cxcl10, Cxcl13, Il2ra, Il2rb, Ccr7*) and tumoricidal factors machinery (*Nos2, Ass1, Prf1, Gzmb*) (Fig. 5C). Downregulated genes functionally belonged to GO terms related mainly to translation and cell division (Fig. 5B), which was in line with a significant decrease of Ki67^+^CD11b^+^ proliferating GAMs upon the RGD+aPD-1 treatment (Fig. 5D). Downregulated genes included *Gpnmb*, coding for transmembrane glycoprotein nonmetastatic B (GPNMB) overexpressed in GAMs in humans and mice, and associated with a poor patient prognosis (41, 42). GPNMB produced by macrophages plays a crucial role in the proneural-mesenchymal transition through immune cell-tumor interplay and GPNMB-high macrophages impair T cell activation by DCs (43). Also, genes involved in mitochondrial electron transport and oxidative phosphorylation were downregulated, indicating a metabolic switch toward proinflammatory myeloid cells (Fig. 5B-C). All those data are consistent with the reduced protumoral activation of GAMs and their transition to the proinflammatory state.

**Fig. 5.**
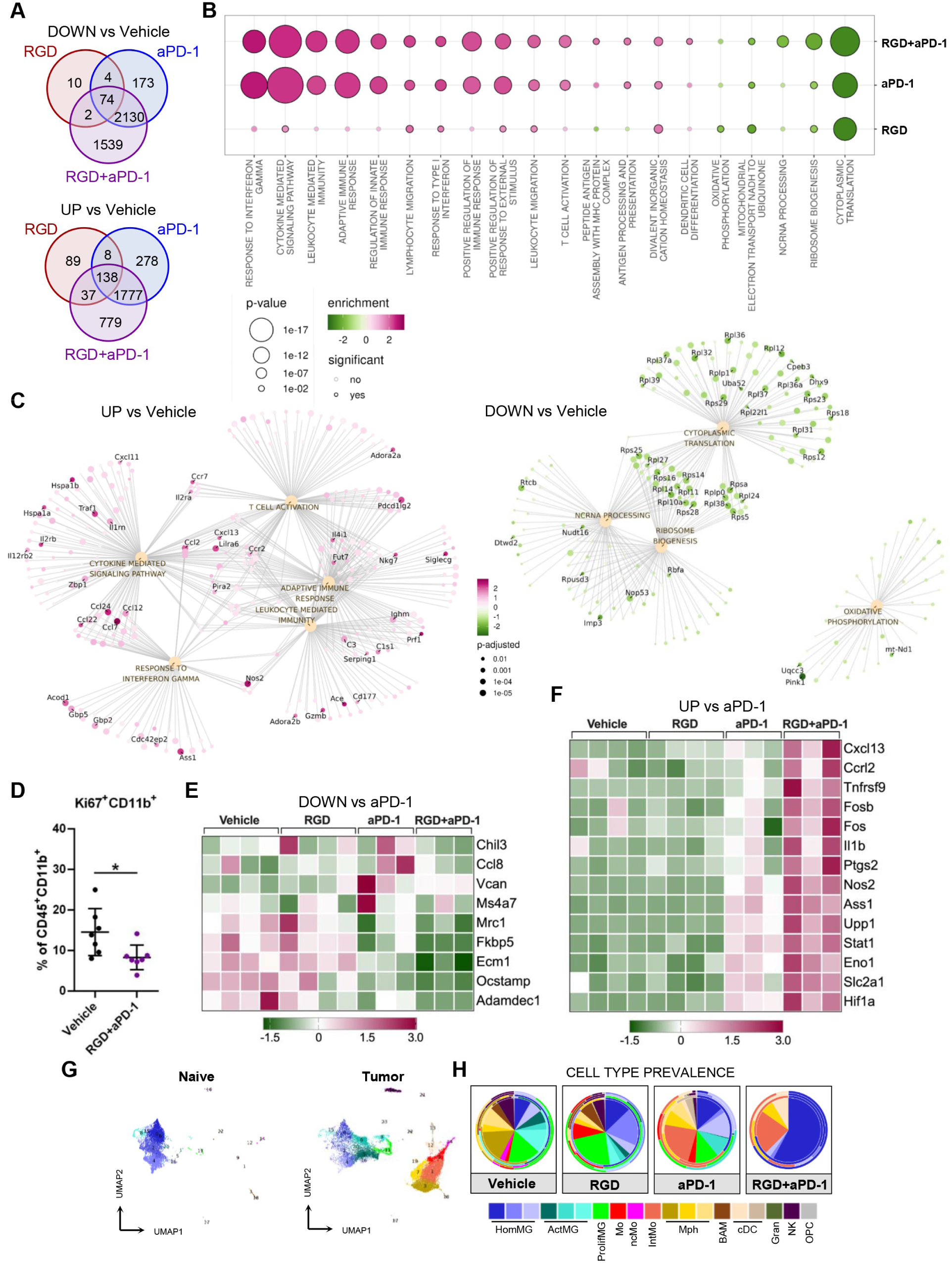
Transcriptomic profiles of CD11b+ from RGD+anti-PD-1 treated mice show reduced expression of the tumor supportive genes and upregulation of antitumor response genes. Gene expression profiling using RNA-seq of CD11b^+^ cells from the tumor-bearing hemispheres of mice from all experimental groups at 28 DPI. (A) Venn diagrams showing a number of down- and upregulated DEGs in RGD, anti-PD-1 (aPD-1) and RGD+aPD-1 groups compared to Vehicle. (B) Functional enrichment of GO biological processes for up- and downregulated genes in RGD+aPD-1 compared to Vehicle. (C) Graphical representation of selected overrepresented GO terms among DEGs in RGD+aPD-1 versus the Vehicle group. Dots represent genes, lines - membership in the given term; top 40 genes with outstanding fold-change and *p*-value were labeled. (D) Percentages of proliferating (Ki67^+^) myeloid cells isolated from the brains of Vehicle and RGD+aPD-1 treated mice. Statistical analysis by Mann-Whitney test; *p < 0.05. (E,F) Z-score heatmaps representing the relative gene expression in CD11b^+^ cells isolated from brains of glioma-bearing mice. Selected down- (E) and upregulated genes (F) comprise the top DEGs in RGD+aPD-1 vs aPD-1. (G-H) Cell type prevalence inferred from deconvolution of bulk RNA-seq data of CD11b^+^ cells sorted at 28 DPI. Deconvolution was based on transcriptomic signatures (see Fig. S4) of myeloid cell clusters form naïve and glioma-bearing brains (G). Predicted cell type abundance is presented on pie-charts (H) with average values in the center and cell prevalence in each repetition indicated on the external rims.

Transcriptomic responses in tumor-infiltrating CD11b^+^ cells from aPD-1 alone and aPD-1+RGD groups were overlapping to a great extent. However, the downregulated genes in the RGD+aPD-1 versus aPD-1 group included known markers of a proinvasive phenotype and suppressors of a proinflammatory switch in macrophages, such as *Chil3*, *Mrc1*, *Fkbp5*, *Ms4a7*, *Vcan*, *Ecm1*, *Ccl8* and *Ocstamp* (Fig. 5E). Among the genes significantly upregulated in RGD+aPD-1 vs aPD-1 we identified: *Ptgs2, Nos2, Ass1, Il1b*, *Stat1* (encoding mediators and enzymes vital for macrophage proinflammatory immune response (44), *Cxcl13* (coding for B cell chemoattractant), *Ccrl2* (a predictive indicator of robust antitumor T-cell responses, selectively expressed in TAMs with the immunostimulatory phenotype in humans and mice (45), *Slc2a1* and *Upp1* (related to glycolysis) and *Tnfrsf9* (coding for CD137/4-1BB), which expression on human monocytes/macrophages leads to their metabolic and functional reprogramming and enhances tumoricidal activity (46) (Fig. 5F). These transcriptomic changes in glioma-associated CD11b^+^ cells delineate the superior efficacy of the RGD+aPD-1 versus aPD-1 treatment.

Finally, through bulk RNA-seq deconvolution (leveraging above mentioned CITEseq data (34), we inferred a cell type abundance within immunosorted glioma-associated CD11b^+^ cells from animals in each group (Fig. 5G-H, Fig. S7). We found significantly higher prevalence of homeostatic microglia and reduced abundance of activated microglia within CD11b^+^ cells in the RGD+aPD-1 treated mice as compared to the vehicle group. Intermediate monocytes and DC signatures were enriched in aPD-1 and RGD+aPD-1 groups, while decreased macrophage scores characterized the response to RGD and RGD+aPD-1. CD11b^+^ cells from the RGD+aPD-1 group showed the predominance of homeostatic microglia signatures, indicative of the phenotype observed in a healthy mouse brain, which corroborated with the lower tumor burden in these mice.

### Targeting intergin signalling has a translational value

The responses of brain myeloid cells to patient-derived orthotopic xenografts in immunodeficient mice follow similar patterns as in syngeneic experimental glioma models (47). The RGD peptide effectively blocked the invasion of human glioma cells in mouse and human microglia co-cultures (Fig.1). Thus, we tested whether intratumorally administered RGD can modify the myeloid TME in intracranial human gliomas in immunodeficient mice. U87-MG RFP^+^ tumor-bearing mice received the vehicle, RGD or the control peptide RAE via osmotic micropumps (Fig. 6A). RGD did not reduce the tumor growth when compared to the vehicle and RAE treatments (Fig. S8A) and the changes of animal body weight over the course of experiment were in the same range in all groups (Fig. S8B). Also, the percentages of microglia and macrophages in the tumor-bearing hemispheres of mice at 21 DPI were similar in all experimental groups (Fig. S8C). However, gene expression profiling of FACS-sorted CD11b^+^ cells indicated the change of the myeloid cells phenotypes upon the RGD treatment (Fig. 6B-C).

**Fig. 6.**
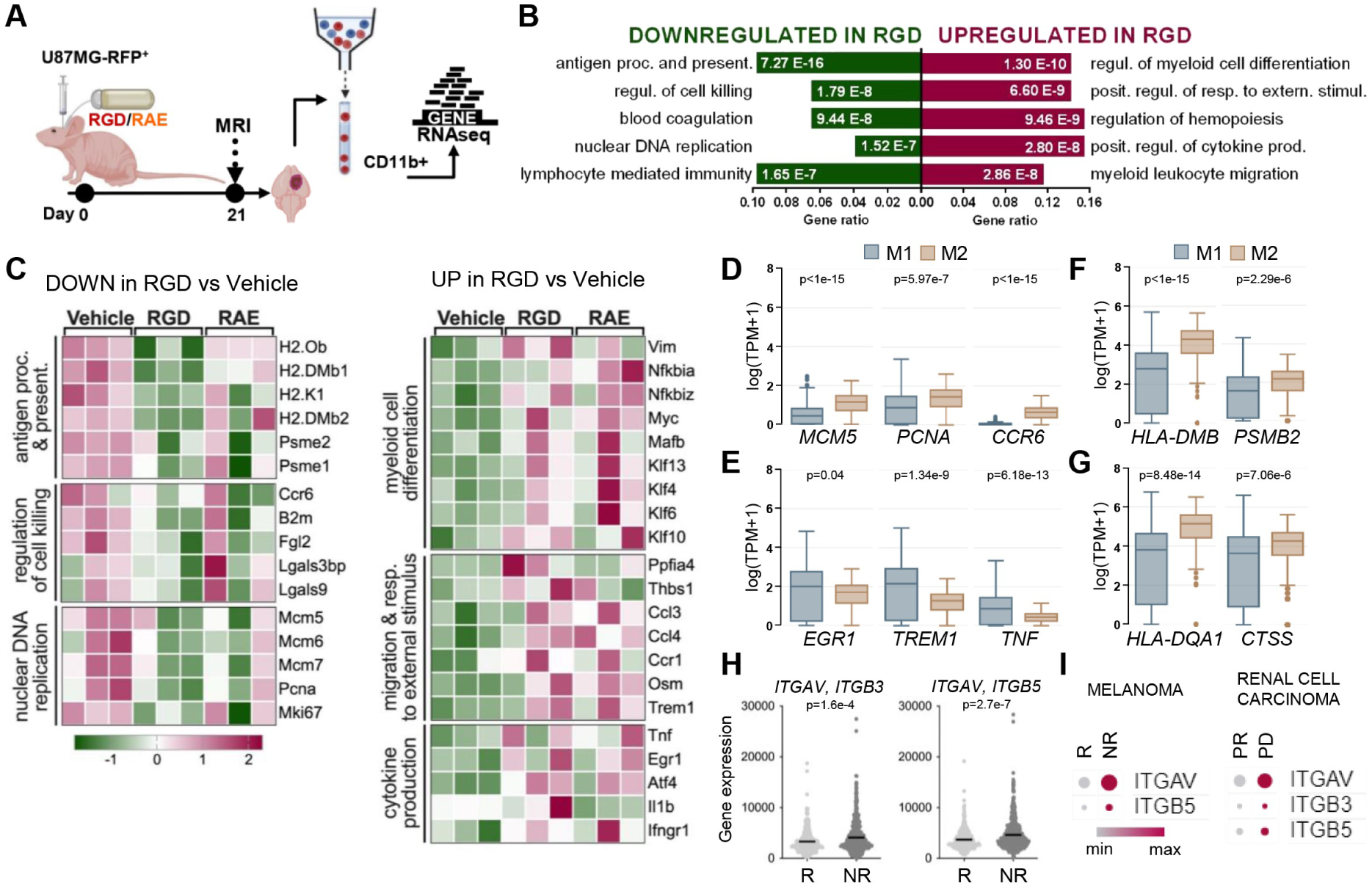
Integrin blockade counteracts the reprogramming of GAMs by human gliomas in mice and emerges as a strategy for improved ICI immunotherapy response. (A) Scheme of the experimental pipeline. Athymic Nude-Foxn1nu mice were orthotopically implanted with U87MG-RFP^+^ human glioma cells and received water (Vehicle), RGD or control peptide (RAE) via osmotic pumps. (B) Functional enrichment analysis with GO biological processes for down- and upregulated genes in CD11b^+^ isolated at 21 DPI from glioma-bearing brain hemispheres of mice treated with RGD as compared to Vehicle. Enriched GO pathways are shown; a size of the bars indicates gene ratios (a number of genes annotated to the pathway/total number of DEGs). (C) Z-score heatmaps with selected up- and downregulated genes in CD11b^+^ cells from RGD vs Vehicle treated animals. (D-G) Differential gene expression in classically (M1) and alternatively (M2) activated macrophages inferred from bulk-RNA-seq data from GBM samples (TCGA) using quanTIseq deconvolution tool (via Gepia2021). Human analogs of the selected genes identified as down- (D, F, G) or upregulated (E) in GAMs after RGD treatment in nude (D, E, F) and C57/Bl6 (G) mice. (H) Levels of *ITGAV/ITGB3* and *ITGAV/ITGB5* in pretreatment tumor biopsies from ICI therapy responders (R) and non-responders (NR) (according to ROCplotter). Statistical analysis by Mann-Whitney test. (I) *ITGAV*, *ITGB3*, *ITGB5* expression in tumor-associated myeloid cells from ICI – treated patients (R – responders, NR – non-responders, PR – partial response, PD – progressive disease). Generated via Single Cell Portal.

GO functional enrichment analysis on DEGs showed that genes involved in antigen processing and presentation, regulation of cell killing, blood coagulation and DNA replication were downregulated in the RGD group as compared to the vehicle (Fig. 6B). Among these genes, we identified minichromosome maintenance protein 5 (*Mcm5*) and its analogs (*Mcm6*, *Mcm7*), proliferating cell nuclear antigen (*Pcna*) and *Ki67*, which indicates the decreased proliferation of GAMs (Fig. 6C). Genes related to antigen processing and presentation, encoding the components of MHC class I (*B2m* and *H2-K1*) and MHC class II (*H2-Ob, H2-DMb1, H2-DMb2)* as well as genes implicated in regulation of cell killing, such as chemokine receptor 6 (*Ccr6*) expressed on immature dendritic cells, galectin-3 binding protein - a negative regulator of NF-κB activation (*Lgals3bp)*, and galectin-9 (*Lgals9)* contributing to M2-type macrophage polarization, were downregulated in the RGD group (Fig. 6C).

The upregulated genes in glioma-associated CD11b^+^ cells from the RGD-treated mice were related to myeloid cell differentiation, response to external stimulus, myeloid leukocyte migration and cytokine production (Fig. 6B). We found high expression of *Il1b*, *Tnf* cytokine coding genes and *Ifngr1*, which encodes the ligand-binding chain of the interferon-gamma receptor. Genes related to myeloid differentiation and linked to macrophage activation and polarization, included *Vim* (induced in TNFα-activated macrophages), *Myc*, *Mafb*, *Nfkbia*, *Nfkbiz*, as well as genes encoding several Kruppel-like factors (*Klf4*, *Klf6*, *Klf10*, *Klf13*) (48), indicating reprogramming in CD11b^+^ cells from RGD treated mice. Interestingly, the RGD treatment induced the expression of genes involved in response to external stimulus, including oncostatin M (*Osm*), which is released by proinflammatory macrophages and has antitumor potential (49) and genes such as *Ccl3, Ccl4* and *Ccr1*, critical for the recruitment of leukocytes to the site of inflammation, and *Trem1* encoding triggering receptor expressed on myeloid cells-1 (TREM1) - a proinflammatory receptor that amplifies antitumor immune responses (50) (Fig. 6C).

In GBM samples from the TCGA database, human analogs of genes downregulated in CD11b^+^ in response to RGD in U87-MG-RFP^+^ gliomas (including genes involved in antigen presentation in both experimental models) were inferred as expressed in alternatively (M2) activated macrophages (Fig. 6D,F-G) and those with upregulated expression in the RGD group were predominantly associated with classically (M1) activated macrophages (Fig. 6E), as predicted using quanTIseq deconvolution of bulk RNA-seq data via GEPIA2021. These data indicate the efficacy of RGD to block the protumoral myeloid cell reprogramming induced by human glioma cells and to induce a re-emergence of macrophages with the antitumor phenotype.

According to analysis leveraging ROC Plotter (https://www.rocplot.com/immune, (51)), *ITGAV*/*ITGB3* and *ITGAV*/*ITGB5* signatures in pretreatment biopsies were significantly higher in non-responders than in responders to ICI immunotherapy in various types of cancers (Fig. 6H). Moreover, exploration of scRNA-seq data (via Single Cell Portal) revealed that the elevated levels of individual integrins in myeloid cells are negative predictors of ICI therapy response in melanoma and renal cell carcinoma patients (Fig. 6I). This suggests that increased expression of these RGD-binding integrins confers resistance to immunotherapies.

## DISCUSSION

Patients with GBM rarely respond to ICIs, with clinical benefits reported in less than 10% of patients, and only in a neoadjuvant setting (8, 52, 53). In this study, we proposed a new approach of anti-glioma therapy based on combining an integrin blockade with PD-1 blockade. We demonstrate strong evidence that the synthetic RGD peptide targeting the integrin-mediated communication between tumor and myeloid cells blocks tumor-induced reprogramming of these cells *in vitro* and *in vivo*. The intratumoral administration of RGD results in converting a “cold” TME to “hot” and stabilization of neoangiogenesis, which boosts innate and adaptive immune responses elicited by anti-PD-1 antibody, leading to augmented antitumor activity.

Despite the well described role of integrins in cancer, their engagement in the functional regulation of tumor-infiltrating myeloid cells has been underexplored. GAMs are the major immune component of GBM, they contribute to tumor progression and impair antitumor responses (6, 7, 15). Preclinical data from rodent glioma models, including ours, point to the importance of αvβ3/αvβ5 integrin signaling in recruitment (24) and phenotype decision making in GAMs (20, 23). Herein, we provide a compelling evidence that RGD - the integrin blocking peptide hinders glioma-microglia interactions inhibiting microglia-dependent invasion of human and mouse glioma cells *in vitro* and prevents the protumorigenic reprogramming of GAMs *in vivo*. Through transcriptome profiling we demonstrate that integrin blockade using RGD induced the phenotype switch of GAMs in GL261/C57BL5 and U87-MG-RFP/ Nude-Foxn1nu mouse models, corroborating our previous data with SPP1 or MFG-E8 knockdown in rats (20). Hallmark protumorigenic genes (including *Arg1*, *Mrc1*, *Il18bp*) were downregulated and genes linked to T cell chemotaxis, cytotoxic NK function and proinflammatory cytokine production were upregulated in GAMs from RGD-treated mice. In line with the transcriptomic data, GL261 tumors in RGD-treated mice were significantly less infiltrated by ARG1^+^ myeloid cells as compared to controls. Arginase deprives arginine that is auxotrophic for T cells. Profound decrease of the frequency of ARG1^+^ GAMs mitigates one of the major obstacles for T cell survival in TME. In our recent study, arginase inhibition combined with anti-PD-1 antibody effectively reduced glioma growth in mice (*26*), prompting us to hypothesize that RGD-driven phenotype switch of GAMs may also contribute to improved response to ICIs and potentially to other immunotherapies.

Due to an intricate relationship between the innate and adaptive immune system, the response of GAMs to RGD treatment was more pronounced in immunocompetent mice than in mice devoid of the adaptive immune system. Interestingly, the top common downregulated pathways in GAMs sorted from RGD-treated mice in both models, were related to antigen processing and presentation. Although this function is a prerequisite for antitumor immune response, in the absence or malfunction of co-stimulatory molecules the result of antigen presentation can render T cells anergic. Glioma-infiltrating microglia/macrophages from postoperative tissue specimens of glioma patients had surface MHC-II expression but lacked expression of the co-stimulatory molecules CD86, CD80, and CD40 critical for T-cell activation (54). MHC-II-expressing blood derived myeloid cells in GL261 tumors in mice displayed a bona fide immunosuppressive and reduced co-stimulatory phenotype (55). Whether RGD-dependent decrease of antigen presentation in immunosuppressed TME may play some role in subsequent response to immunotherapy requires further studies.

The αvβ3 and αvβ5 integrin receptors are strongly overexpressed on the proangiogenic endothelial cells, including those forming tumor-associated blood vessels in GBM and inhibitors targeting these integrins, such as a cyclic RGD peptide, cilengitide (CIL), act as anti-angiogenic agents (56). We report that the intratumoral delivery of RGD resulted in improved vessel density. It corroborates the paradoxical proangiogenic effect reported with CIL (at low doses), which led to the vascular normalization, i.e. remodeling of the structurally and functionally immature tumor vasculature resulting in improved blood flow and drug delivery (57). This phenomenon might contribute to improved antitumor effects of radiotherapy or chemotherapy upon co-administration with CIL (58, 59) and the clinical benefit apparently seen in some GBM patients with MGMT promoter methylation in the Phase III clinical trial with CIL added to the standard chemoradiotherapy (60). Vascular normalization transiently induced by anti-angiogenic therapy promotes the infiltration of T lymphocytes, diminishes hypoxia and thereby immunosuppression within the tumors (61–63). Anti-VEGF drugs enhanced the efficacy of ICIs in preclinical settings, also in GBM models (16, 17). Hence, increased density of structurally improved blood vessels observed upon the intratumoral delivery of RGD might have contributed to the boosted response to anti-PD-1 immunotherapy in glioma-bearing mice.

Profiling the immune landscapes of human and mouse tumors identified the scarcity of activated cytotoxic T cells, plenitude of suppressive myeloid cell subsets and poor antigen presentation in the brain as the major obstacles for effective immunotherapy of GBM (10, 40). According to a recent single-cell study, neoadjuvant PD-1 blockade induced T cell and cDC1 activation but failed to overcome the immunosuppressive GAMs in recurrent human GBM (64). Therefore, several combination therapies of ICIs have been tested in preclinical settings with blocking the infiltration or re-educating suppressive myeloid cells (9, 40, 65). In the current study, we showed the synergy between the intratumorally-delivered RGD and systemic administration of anti-PD-1 in mice harboring syngeneic GL261 tumors. Anti-PD-1 treatment either alone or in combination led to decrease of protumoral marker genes (such as *Gpnmb* and *Mrc1*) and upregulated expression of genes related to antigen presentation, interferon signaling, innate immune response and secretion of T cell chemotactic factors. However, ICI monotherapy induced the previously reported negative feedback response (66) characterized by the upregulation of ARG1 and PD-L1 on myeloid cells and sustained levels of Treg cells, counteracting the re-activation of T cells after immunotherapy. PD-L1 expression on macrophages has been associated with poor survival and resistance to immunotherapy in GBM patients (40, 67). Breaking the feedback response and reduced expression of the immunosuppressive molecules in GAMs upon co-administration of RGD likely contributed to the efficacy of the combined treatment. Accordingly, the transcriptomes of glioma-associated CD11b^+^ cells from aPD-1 and RGD+aPD-1-treated mice indicated a complete phenotypic switch only in the latter group. Intratumoral microglia and more importantly blood-derived macrophages/DCs from RGD+aPD-1 mice showed elevated levels of MHC-II. MHC-II-restricted antigen presentation by cells of the myeloid compartment, particularly those infiltrating from the periphery but not microglia alone, is required for response to ICI and correlates with favorable clinical outcome of GBM patients undergoing anti-PD-1 therapy (55).

Reprogramming of GAMs in RGD+aPD-1 treated mice was associated with improved T cell immune surveillance within the TME, reflected by increased frequency and proliferation of intratumoral CD8^+^ T cells, reduced numbers of Tregs and higher ratios of CD8^+^/Treg and CD8^+^/CD11b^+^. Moreover, RGD+aPD-1 treatment instigated the IFNγ production by tumor-infiltrating T cells, including CD8^+^ effector T cells, and elevated its intratumoral levels. Consistent with the augmented antitumor response, the highest concentration of cytolytic granzyme B and TNFα were detected in the immunostimulatory milieu of RGD+aPD-1-treated tumors, which can attract additional T cells and professional antigen presenting cells (e.g., dendritic cells) by means of elevated CCL3, CCL5 and CXCL10. Thus, the combined treatment reshaped the glioma immune landscape and incapacitated several mechanisms dampening the proper action of T cells in TME, resulting in enhanced T cell proliferation and activation needed for functional antitumor responses. GBM remains an incurable disease and improving the treatment remains a paramount challenge for clinicians and researchers. Here we show the synergy between two therapeutic modalities that have failed individually: an immune check-point and integrin blockade. We reveal that the RGD peptide modifies the function of GAMs and tumor neovasculature. Bimodal action of the peptide reshapes the glioma TME and facilitates the successful revival of antitumor host response with ICI. While the results show new therapeutic opportunities for GBM patients, some limitations to our work should be considered. Synergistic effects of ICI and RGD could not be tested in immunodeficient mice, however we show the efficacy of RGD in blocking the reprogramming of myeloid cells in human U87-MG gliomas *in vivo* and in co-culture models. RGD was delivered throughout the experiment accomplishing the preconditioning of immune-suppressed TME via integrin antagonism. Investigations into an optimal duration of treatment and the best timing of administration of PD-1 inhibitors may further enhance the efficacy of the combinatorial approach. A potential caveat of our strategy comes from the limited peptide stability upon systemic administration. Development of modified peptides or convection-enhanced delivery could overcome such challenges in the future.

Integrin-coding genes are evidently upregulated in GBM, expressed on myeloid cells, pericytes and endothelial cells, and their high expression is associated with worse outcome for patients (Fig. 1B,C) and higher risk of ICI therapy failure (Fig. 6 H,I). Therefore, αvβ3/5-integrins are valid therapeutic targets in GBM. The successful use of integrin-blocking agents requires proper patient stratification based on the ligand (eg. SPP1) or target integrin expression. In a retrospective study, higher αvβ3 protein levels were associated with improved survival of GBM patients treated with CIL (68). The presented results encourage further studies of integrin signaling inhibition in human GBM and indicate its potential clinical application. Finally, re-investigation of patient responses to previous anti-integrin therapies from the perspective of their impact on myeloid cells may shed light on potential benefits from combination of integrin blockers with immunotherapy.

## METHODS

### Sex as a biological variable

Experiments were perfored on male mice because male animals exhibited less variability in tumor growth.

### Cell cultures

Murine glioma GL261 cells were obtained from Prof. Helmut Kettenman (MDC, Berlin, Germany) and modified to GL261 tdTomato+luc+ glioma cells as described (15). Murine and human glioma cell lines: U-251 MG, U87-MG (ATCC, Manassas, VA) and U87-MG RFP+ (AntiCancer Inc., San Diego, CA, USA) were cultured in Dulbecco’s modified Eagle’s medium (DMEM) with 10% fetal bovine serum (FBS) (Gibco, MD,USA). After thawing, GL261 tdTomato+luc + and U87-MG RFP+ cells were supplemented with 400 μg/ml G418 (Roche, Manheim, Germany) for two passages. Human SV40 immortalized microglial cells (HMSV40) were cultured in PriCoat T25 flasks in Prigrow III Medium (Applied Biological Materials, Richmond, Canada) supplemented with 10% FBS (Gibco, MD, USA). Murine immortalized microglial BV2 cells (received from Prof. Klaus Reymann from Leibniz Institute for Neurobiology) were cultured in DMEM GlutaMAX™ supplemented with 2% FBS (Gibco, MD, USA). All cells were cultured with antibiotics (100 U/ml penicillin,100 µg/ml streptomycin) in a humidified atmosphere of CO_2_ /air (5%/95%) at 37°C (Heraeus, Hanau, Germany). Mycoplasma contamination-free status was checked regularly.

### Cell viability and proliferation assays

Cell viability was evaluated using MTT metabolism test as described [Ciechomska et al. 2023]. Cell proliferation was measured using BrdU ELISA kit (Roche, Mannheim, Germany) according to the manufacturer’s protocol. Briefly, 5 × 10^3^ cells were seeded onto 96-well plates. The next day the media was exchanged to DMEM GlutaMAX™ supplemented with 2% FBS (Gibco, MD, USA) and RGD and RAE peptides were added at a final concentration 100 μM for 18 h. Peptides with N terminal acetylation and C terminal amidation were purchased from Genscript (Rijswijk, Netherlands). Peptides were dissolved in DMSO, the 0.2% solvent was added as a control. MTT solution (0.5 mg/mL; Sigma-Aldrich, Taufkirchen, Germany) or BrdU were added for 1.5 and 2 h, respectively. Optical densities were measured at 570 nm for MTT and 450 nm for BrdU assays, using a scanning multi-well spectrophotometer.

### Invasion assays

Invasion assay was performed as described (*26*). BV2 and HMSV40 cells (4×10^4^) were plated onto a 24-well plate and co-cultured with GL261 cells (1×10^5^/insert), U87 and U251 (4×10^4^/insert) on Matrigel-covered membranes in 2% FBS containing media. Cells were treated with 100 μM RGD and RAE (control) peptides or 0.2% DMSO. GL261 were co-cultured with BV2 cells for 24 h and human glioma cells with BV2 or HMSV40 cells for 18 h.

### Peptide stability

The RGD peptide was dissolved in water at a concentration of 2 mg/ml and was incubated in osmotic pumps for 1, 7 and 14 days at 37°C. The peptide concentration was measured using Jupiter Proteo C12 2.1 x 250 mm, 4 μm column (Phenomenex, Torrance, USA) and HPLC Prominence coupled LCMS-IT-TOF (Shimadzu, Duisburg, Germany) under the following conditions: Phase A: 0.1% HCOOH in MilliQ water, phase B: 0.1% HCOOH in ACN, gradient conditions: 0 min 2% B, 20 min 50% B, 25 min 95% B, 30 min 95% B, 35 min 5% B, 55 min 5% B at a 0.2 ml/min flow rate. 1 ml of sample was injected. A Shimadzu IT-TOF ESI-MS system was used for the mass analysis in the automatic mode setting with a scan range of 150 - 2000 Da and an ion accumulation time of 10 ms. Electrospray ionization was performed in the positive ionization mode with a spray capillary voltage of 5.0 kV. The interface temperature was kept at 250°C and heat block at 520°C. Nebulizing gas was introduced at 1.5 L/min, while the drying gas was set to 100 kPa. Mass spectra were recorded in positive ion mode using the LCMSsolution software provided by Shimadzu. The analysis was performed at the Department of Chemistry of the Warsaw University, Poland.

### Stereotactic implantation of glioma cells and treatments

Male C57BL/6 mice or Athymic Nude-Foxn1nu mice (Charles River Laboratories, USA) 10-12 weeks old were housed with free access to food and water, on a 12 h/12 h day and night cycle. All efforts have been made to minimize the number of animals and animal suffering. Mice were anesthetized with isoflurane (4–5% induction, 1–2% maintenance) using Isoflurane vaporiser (Temsega, Tabletop Anesthesia Station). Before starting the surgical procedure and during the surgery a depth of anesthesia was verified. Choice of anesthetics was recommended by the veterinarian and approved by The Local Ethics Committee. GL261 tdTomato^+^luc^+^ glioma cells (80,000 in 1 μL of DMEM) in C57BL/6J or U87-MG RFP (50,000 in 1 μl of DMEM) in Athymic Nude-Fox1nu mice were stereotactically injected to the right striatum at the following coordinates: 1 mm anterior and 2 mm lateral from bregma, 3 mm deep from the surface of the brain. At the same time, Alzet osmotic micropumps (DURECT Corporation, Cupertino, CA, USA) were installed in a subcutaneous pocket on the back, slightly posterior to the scapulae. A small incision was made in the shaved skin, and a hemostat was used to create the subcutaneous pocket for the pump which was then inserted into the pocket and the wound was closed with tissue glue. Prior to implantation, the metal flow moderators were replaced with PEEK flow moderators to be MRI compatible. To ensure that the pumps were active when implanted, the filled pumps were placed in sterile saline at 37°C for 24 h before implantation. Osmotic pumps were filled with: H_2_O (Vehicle), RGD or RAE peptides at a concentration of 2 mg/ml in H_2_O and by means of a catheter they continuously delivered the solutions intratumorally at a controlled rate of 0.11 μL/h for the duration of 21 or 28 days.

The mice were monitored until they completely recovered from anesthesia. The animals were weighed weekly and observed daily for clinical symptoms and evidence of toxicity by evaluating their eating, mobility, weight loss, hair loss, and hunched posture. Anti-PD-1 antibody (Biolegend, GoInVivo™ Purified anti-mouse CD279) was injected intraperitoneally (i.p.) at a dose of 10 mg/kg on days 8, 10, 12, 14 post-implantation. Control groups received IgG antibody by i.p. injection. Animals were euthanized when they lost more than 20% of body weight compared to day 0.

### Tumor size measurement using magnetic resonance imaging

Heads of the animals were scanned with 7T BioSpec 70/30 MR system (Bruker, Ettlingen, Germany) equipped with Avance III console and actively shielded gradient system B-GA 20S (amplitude 200 mT/m) with an integrated shim set up to 2^nd^ order. A combination of transmit cylindrical radiofrequency volume coil (8.6 cm inner diameter, Bruker) and head-mounted mouse dedicated receive-only array coil (2×2 elements, Bruker) was used. The animals were anesthetized at 1.5-2% isoflurane (Baxter, Deerfield, IL, USA) in oxygen and positioned prone with the head placed in the MR-compatible bed integrated with an anesthesia mask. Respiration and rectal temperature were monitored throughout the experiment with a MR-compatible small animal monitoring system (SA Instruments, Stony Brook, NY, USA). All the imaging sessions started with a localizer protocol consisting of three orthogonal scout scans to accurately position the animal inside the magnet center. To evaluate volumes of the brain structures, structural transverse MR images covering the whole brain were acquired with T2-weighted TurboRARE (TR/TEeff = 7000/30ms, RARE factor = 4, spatial resolution = 86μm x 86μm x 350μm, 42 slices, no gaps, number of averages (NA) = 4, scan time ∼ 23min). MRI scans were evaluated using OsiriX software (Pixmeo, Geneve, Switzerland) and manually delineated tumor regions on image series were used for volumetric assessments.

### Cytokine analysis

Pro- and anti-inflammatory cytokines were measured in serum and brain homogenates from control and treated animals. Blood was collected before perfusion and allowed to clot for 30 min before centrifugation (10,000 × g, 10 min at room temperature). The serum was collected and stored at - 80°C. Brain homogenates were prepared by adding an equal volume of Cell Lysis Buffer 2 (R&D Systems, Minneapolis, MN, USA) to dissociated brain tissues. Samples were then incubated at room temperature for 30 min with gentle agitation and debris were removed by centrifugation. The levels of cytokines were measured using the Luminex Assay Mouse Premixed Multi-Analyte Kit (R&D Systems, Minneapolis, MN, USA) according to the protocol. Cytokine levels were determined using the MAGPIX Multiplexing Instrument (Luminex, TX, USA) with XPonent software. Results for each cytokine were expressed as pg/ml for serum samples and pg/mg of protein for brain homogenates. Protein concentration in brain homogenates was estimated using Bradford reagent (SigmaAldrich) and optical densities at 570 nm were measured using a scanning multi-well spectrophotometer.

### Immunohistochemistry on brain slices

The animals were sacrificed 21 days after GL261 tdTomato^+^luc^+^ cell implantation and perfused with 4% paraformaldehyde in phosphate-buffered saline (PBS). Brains were removed, post-fixed for 48 h in the same fixative solution and placed in 30% sucrose in PBS at 4°C until the tissue sunk to the bottom of the flask. Tissue was frozen in Tissue Freezing Medium (Leica Biosystems, Richmond, IL, USA) and cut in 12 µm coronal sections using a cryostat. The slides were dried at room temperature for 2 h after being transferred from the −80°C storage. Cryosections were blocked in PBS containing 10% donkey serum and 0.1% Triton X-100 for 2 h and incubated overnight at 4°C with rabbit anti-Iba-1 and goat anti-Arg1 antibodies or FITC-conjugated *Lycopersicon esculentum* (Tomato) Lectin (Vector Labs, FL-1171-1). Next, sections were washed in PBS and incubated with corresponding secondary antibodies for 2 h at room temperature. All antibodies were diluted in 0.1% Triton X-100/PBS solution containing 3% donkey serum. Nuclei were counterstained with DAPI (0.001 mg/ml). Images were acquired using the Olympus microscope (Fluoview, FV10i) and Leica DM4000B fluorescent microscope. For reagent specifications, catalogue numbers, and concentrations, see the Supplementary Table 1. We quantified % of Iba^+^ Arg1^+^ cells in 3 different areas of the tumor per each animal. For lectin staining three different sections from each mouse brain were analyzed. In each section 2 regions of interest (ROIs) in the ipsilateral side (near the tumor core) were analyzed.

### Tissue dissociation, flow cytometry and FACS sorting

On day 21 or 28 after GL261 tdTomato^+^luc^+^ or U87-MG RFP+ cell implantation mice were perfused transcardially with cold PBS prior to excision of the brain and spleen. The tumor-bearing hemispheres were dissociated enzymatically with Collagenase type IV and DNase I (both from Merck, Darmstadt, Germany) at a final concentration of 2.5 mg/ml and 0.5 mg/ml, respectively, using gentleMACS Octo Dissociator (Miltenyi Biotec), according to the manufacturer’s protocol.

Next, the enzymatic reaction was stopped by the addition of Hank’s Balanced Salt Solution with calcium and magnesium (Gibco, Germany). The resulting single cell suspension was filtered through 70 μm and 40 μm strainers, and centrifuged at 300 × g, 4°C for 10 min. Myelin was removed by density gradient centrifugation in 22% Percoll as described (27). Next, cells were collected, washed with PBS and counted using NucleoCounter (Chemometec, Gydevang, Denmark).

The spleen was passed through a 70 μm strainer and gently grounded using the plunger of a syringe with 3 ml of PBS to yield a single cell suspension. Following centrifugation (300 x g, 5 min), red blood cells were lysed using ACK Lysing Buffer (Life Technologies, Grand Island, NY, USA) for 10 min at room temperature. The splenocytes were collected by centrifugation, washed with PBS and counted using NucleoCounter (Chemometec, Gydevang, Denmark).

For preparation to flow cytometry analysis samples were handled on ice and protected from light exposure. Prior to staining with antibodies, samples were incubated with eFluor 506 fixable viability dye (ThermoFisher) in PBS for 10 min. Next, samples were incubated for 10 min with rat anti-mouse CD16/CD32 Fc Block™ (BD Pharmingen) in Stain Buffer (BD Pharmingen) to block FcγRIII/II and reduce nonspecific antibody binding. Then, the cell suspensions were incubated for 30 min with an antibody cocktail in Stain Buffer (BD Pharmingen) for detection of surphace antigens. For intracellular staining, cells were fixed and permeabilized (Foxp3 fixation/permeabilization buffer, eBioscience) prior to incubation with antibodies. List of antibodies – see Table S1. For FACS sorting the cells were stained with CD11b (M1/70 clone) antibody labeled with FITC (BD Pharmingen) and CD45 (30-F1 clone) labeled with PE-Cy7 (BD Pharmingen).

For intracellular cytokine staining, freshly isolated cells from the tumor-bearing brain hemispheres were resuspended in stimulating culture media with 50 ng/ml PMA, 1 µg/ml ionomycin (Sigma Aldrich) and protein transport inhibitor cocktail (brefeldin A and monensin at final concentrations of 10.6 µM and 0.2 mM, respectively; Life Technologies, Carlsbad, CA, USA) for 4 h and then processed for staining of surface and intracellular antigens as described above.

All antibodies were titrated prior to staining to establish the amount yielding the best stain index. Data were acquired using a BD LSR Fortessa Analyzer cytometer and analyzed with FlowJo software (v. 10.5.3, FlowJo LLC, BD). Gates were set based on FMO (fluorescence minus one) controls and back-gating analysis. Percentages on cytograms were given as the percentage of a parental gate. For computational analyses, each sample was downsampled to obtain 10000 CD45^+^CD11b^+^ or CD45^+^CD11b^-^. For tSNE all samples were concatenated and processed using specific plugins in FlowJo v10.

CD11b^+^ cells were FACS sorted from naive or tumor-bearing hemispheres (pooled from 2 animals per sample at day 21 and from individual mice at day 28) using Cell Sorter BD FACSAriaII. All flow cytometry experiments were performed at the Laboratory of Cytometry, Nencki Institute of Experimental Biology. For reagent specifications, catalogue numbers and dilutions see the Supplementary Table 1. Gating strategies used in the analysis are shown in the Fig. S9 and Fig. S10.

### RNA isolation, mRNA library preparation and RNA-sequencing

Immediately after sorting, CD11b^+^ cells were centrifuged and lysed for further isolation of RNA using the RNeasy Plus Mini Kit (Qiagen, Germany) according to manufacturer’s protocol. The integrity and quality of RNA were assessed on an Agilent 2100 Bioanalyzer with an RNA 6000 Pico Kit (Agilent Technologies, CA, USA). Strand-specific RNA libraries were prepared for sequencing (3 - 4 biological replicates/treatment) using a KAPA Stranded mRNA-Seq Kit (Kapa Biosystems, MA, USA). Poly-A mRNAs were purified from 100 ng of total RNA using poly-T-oligo-magnetic beads (Kapa Biosystems, MA, USA). mRNAs were fragmented and a first-strand cDNA was synthesized using reverse transcriptase and random hexamers. A second-strand cDNA synthesis was performed by removing RNA templates and synthesizing replacement strands, incorporating dUTP in place of dTTP to generate double-stranded (ds) cDNA. dsDNA was then subjected to addition of “A” bases to the 3′ ends and ligation of adapters from NEB, followed by uracil digestion by USER enzyme (NEB, MA, USA). Amplification of fragments with adapters ligated on both ends was performed by PCR using primers containing TruSeq barcodes (NEB, Ipswich, MA, USA). Final libraries were analyzed using Bioanalyzer and Agilent DNA High Sensitivity chips (Agilent Technologies, Santa Clara, CA, USA) to confirm fragment sizes (∼300 bp). Quantification was performed using a Quantus fluorometer and the QuantiFluor dsDNA System (Promega, Madison, Wisconsin, US). Libraries were loaded onto a rapid run flow cell at a concentration of 8.5 pM onto a rapid run flow cell and sequenced on an Illumina HiSeq 1500 paired-end.

### Data processing and analysis

Illumina-specific adapters, short reads, and low quality 5′ and 3′ bases were filtered out in the FASTQ format files using Trimmomatic [10.1093/bioinformatics/btu170]] tool (version 0.36). The resulting RNA sequencing reads were aligned to a reference mouse genome sequence (mm10) with STAR aligner (69) (version 2.6.1b) using the two pass Mode Basic option. Duplicate reads were then identified and flagged using Picard Tools (version 2.17.1) [broadinstitute.github.io/picard/]. Quantification of mapped reads and summarization by gene was performed using HTSeq-count (version 0.11.1), with paired mode (-p) and reverse stranded mode (-s reverse) enabled, and only reads with MapQ values of 10 or higher were considered. Low-expressed features were filtered out and an analysis of differentially expressed genes was performed using NOIseq. Only mRNAs encoding protein-coding genes were retained for downstream analysis.

To identify transcriptomic differences between groups, differential expression analysis was performed using NOIseq methods, with the control state (Vehicle) as the reference group and compared with the RGD, RAE or RGD, anti-PD-1 and combination of RGD+anti-PD-1. The variance stabilizing transformation (vst function) was used for visualization. Pathway enrichment analysis was performed utilizing the fgsea (70) on genes ranked by the fold-change. Gene Ontology Biological Processes (GO: BP) was used to better understand the mechanistic findings of the enriched gene lists. The clusterProfiler, VennDiagram and ggplot2 R packages were used to visualize that data.

Bulk RNA-seq data deconvolution was performed using the bisque deconvolution tool (71) with default settings. Deconvolution was based on the cell-type specific transcriptomic signatures generated in our laboratory using CITE-seq on isolated brain CD11b^+^ cells derived from the same glioma model in mice (34).

### Statistical analysis

All *in vitro* data represent at least three independent experiments, in duplicates or triplicates. Statistical significance for invasion assay results was calculated using the chi-square test. The numbers of animals per group are specified in figure captions. Comparisons between two groups were performed with a two-tailed student’s t test. In cases of multi-group comparison one way analysis of variance (ANOVA) was used, followed by Tukey’s HSD test. All statistical analyses were performed using GraphPad Prism 7.0 software. *P* < 0.05 were considered as statistically significant.

### Study approval

All animal experiments were carried out in strict accordance with the Guidelines for the Care and Use of Laboratory Animals (European and national regulations 2010/63/UE September 22, 2010 and Dz. Urz. UE L276/20.10.2010, respectively) and were approved (approval no. 812/2019;1049/2020) by The First Warsaw Local Ethics Committee for Animal Experimentation.

## Data availability

All data associated with this study are in the paper and/or the Supplementary Materials. The RNA-seq datasets have been deposited in NCBI’s Gene Expression Omnibus with GSE266110 identifier. The materials generated in this study can be requested from B.K. or A.E.M..

## AUTHOR CONTRIBUTIONS

Conceptualization and design of experiments: AEM, PPK, KP, SC, BK

Methodology: PPK, SC, JS, KP, AEM, ARC, SB, BG

Investigation: PPK, KP, AEM, SC, JS, MG, MP, YH, BG

Data analysis: ARC, SB, KW

Visualization: PPK, AEM, KP, ARC, SB

Funding acquisition and project administration: AEM

Supervision: BK, AEM

Writing - original draft: PPK, AEM, KP, review & editing: AEM, BK

All authors read and edited the manuscript.

## ACKNOWLEDGMENTS

The authors are grateful for financial support from National Science Centre Poland (no. 2017/25/B/NZ3/02483 to AEM). JS is supported by European Molecular Biology Organization (EMBO) postdoctoral fellowship ALTF 663-2022. M.G. was supported by PACIFIC Call 1 (Polish Academy of Science) Agreement No. PAN.BFB.S.BDN.619.022.2021 and the European Union’s Horizon 2020 research and innovation programme under the Marie Skłodowska-Curie grant agreement No 847639 and from the Ministry of Education and Science. We thank Beata Kaza for her technical assistance. Experiments were performed at the Laboratory of Cytometry and Laboratory of Sequencing - Nencki Institute core facilities.

---

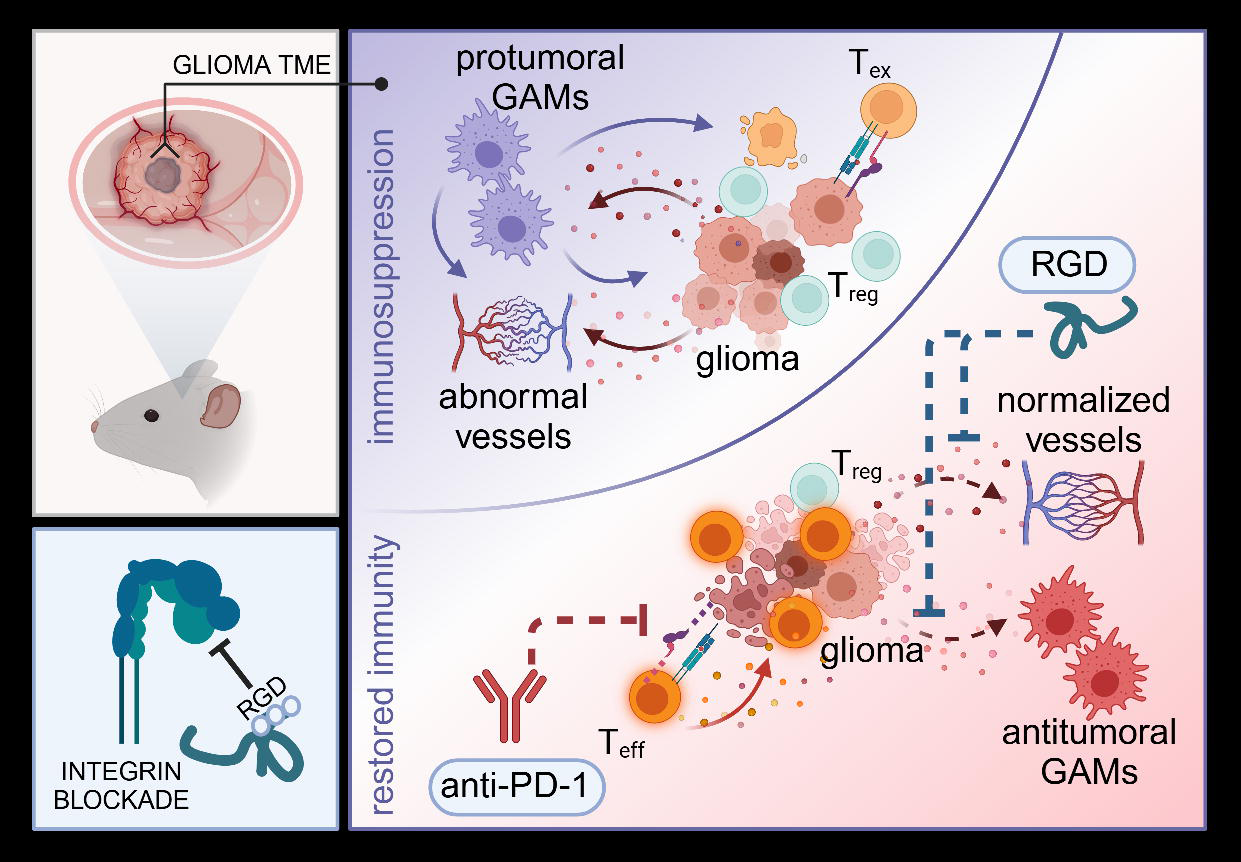

## Notes

**Conflict-of-interest statement**: The authors have declared that no conflict of interest exists.

### Competing Interest Statement

The authors have declared no competing interest.

