## Supplementary Materials for "Integrin blocking peptide reverses immunosuppression in experimental gliomas and improves anti-PD-1 therapy outcome"

**This file includes:**

- Fig. S1. RGD motif-binding integrins in human glioma samples.
- Fig. S2. The RGD peptide does not affect cell viability and proliferation.
- Fig. S3. Stability and biodistribution of the RGD peptide delivered in osmotic pumps.
- Fig. S4. Profiles of myeloid cell in the brain of RGD-treated GL261 glioma-bearing mice.
- Fig. S5. Characterization of the immune microenvironment of GL261 gliomas upon RGD or/and anti-PD-1 treatment.
- Fig. S6. Immune cells phenotyping in glioma TME and spleens at 21 DPI in tumor-bearing mice upon administration of RGD+anti-PD-1.
- Fig. S7. Myeloid cell profiles in the brains of RGD-, anti-PD-1 and RGD+anti-PD-1-treated GL261 glioma-bearing mice.
- Fig. S8. The response to RGD-treatment in U87MG-RFP+ glioma-bearing mice.
- Fig. S9. Gating strategy used for FACS-sorting.
- Fig. S10. Gating strategy used for flow cytometry analysis.
- Table S1. Antibodies.

### Supplementary Fig. 1

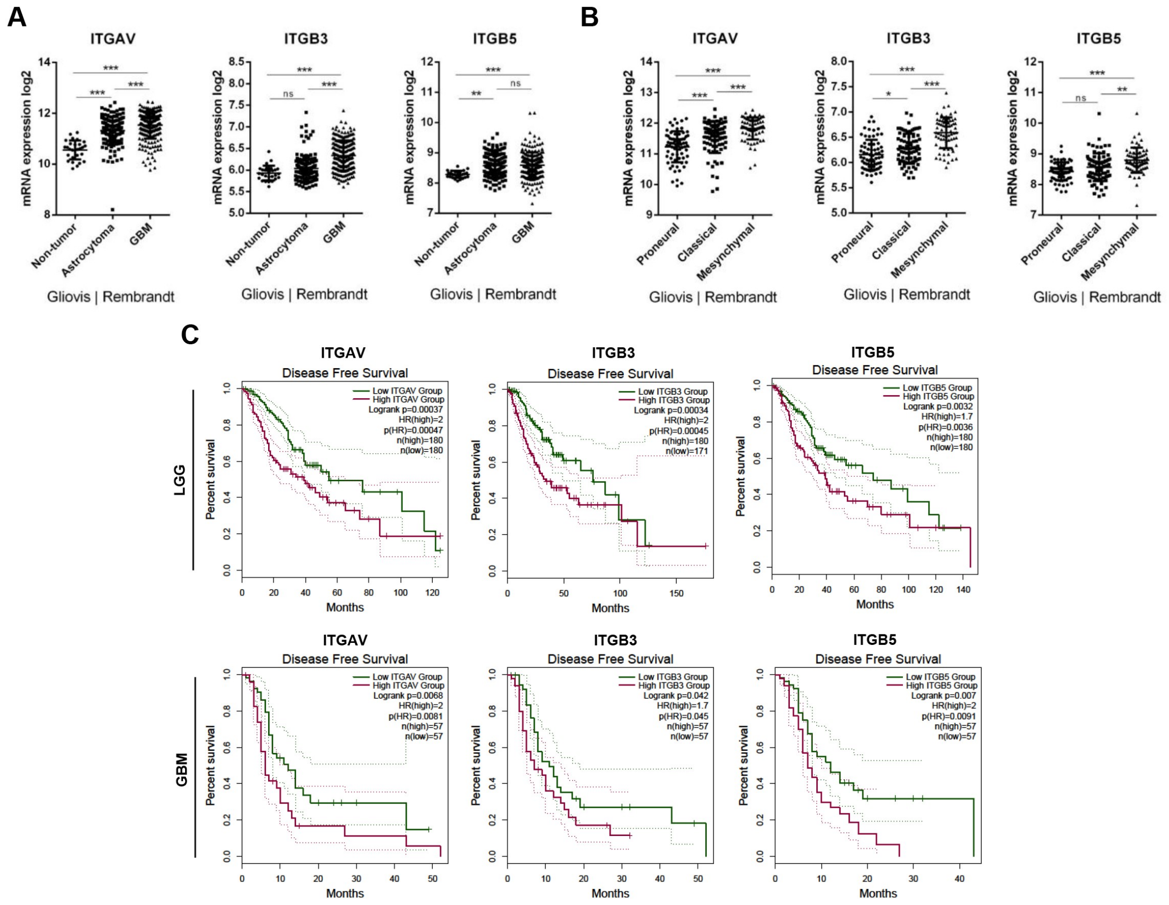


**Fig. S1. RGD-motif binding integrins in human glioma samples.** (A,B) Gene expression data from Rembrandt database extracted using the Gliovis portal (doi: 10.1093/neuonc/now247). (C) Kaplan-Meier curves generated using GEPIA 2 (Gene expression Profiling Interactive Analysis) showing disease free survival of low-grade glioma (LGG, upper panel) and GBM (lower panel) patients with high (in purple) and low (in green) expression status of integrin genes. Samples were grouped with cutoff-high 65% and cutoff-low 35%; dotted line depicts 95% CI.

**Supplementary Fig. 2**

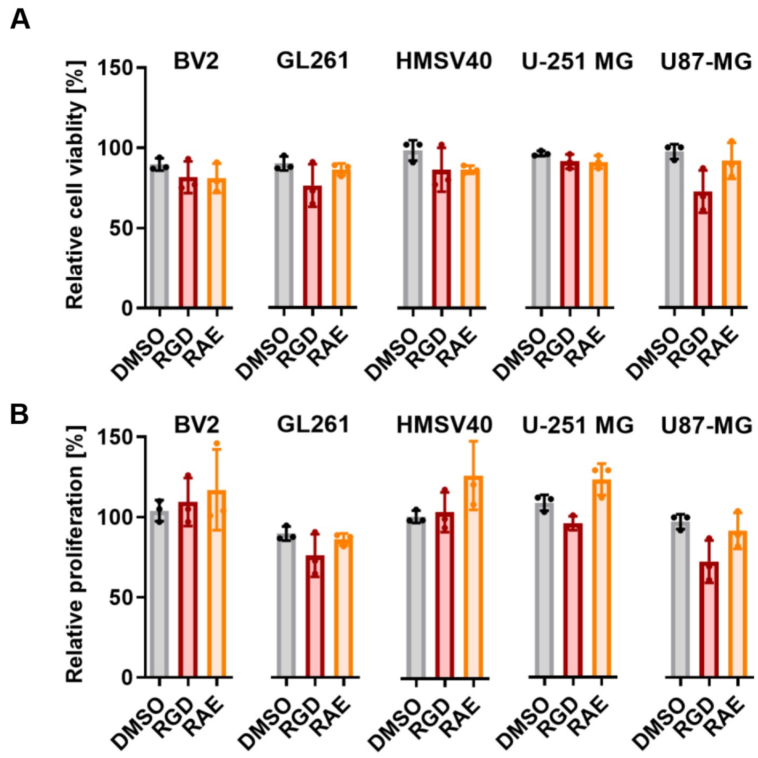


**Fig. S2. The RGD peptide does not affect cell viability and proliferation.** The effects of RGD and RAE peptides on cell viability (A) and proliferation (B) of BV2, GL261, HMSV40, U251-MG and U87-MG cells were determined by MTT metabolism and BrdU incorporation tests. Cell viability and proliferation of untreated cells was set as 100%; n=3.

**Supplementary Fig. 3**

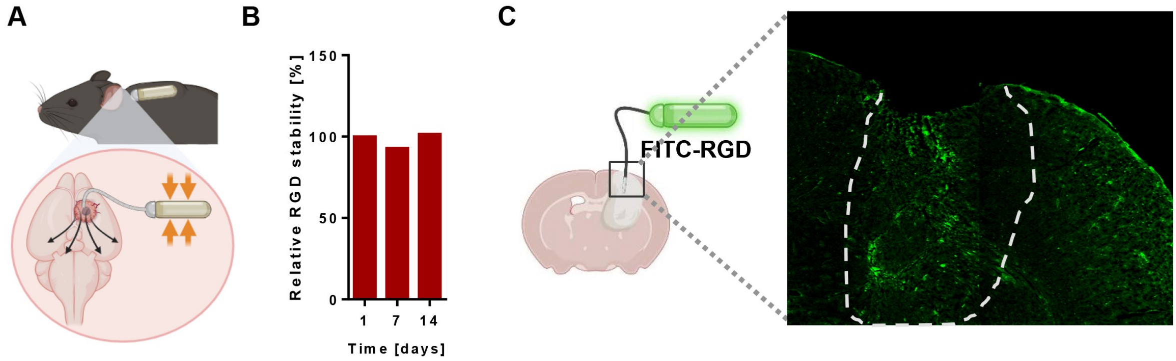


**Fig. S3. Stability and biodistribution of the RGD peptide delivered in osmotic pumps** (A) In-body location and mode of action of osmotic micropumps. (B) Stability of the RGD peptide dissolved in water and incubated in osmotic pumps for 1, 7 and 14 days at 37°C was analyzed using mass spectrometry. (C) Biodistribution of FITC-labeled RGD peptide (green) delivered intratumorally via osmotic pumps to GL261wt tumor-bearing mice. Representative images of the brain section (the ipsilateral hemisphere) at day 14 DPI; the tumor area depicted with a dashed line.

**Supplementary Fig. 4**

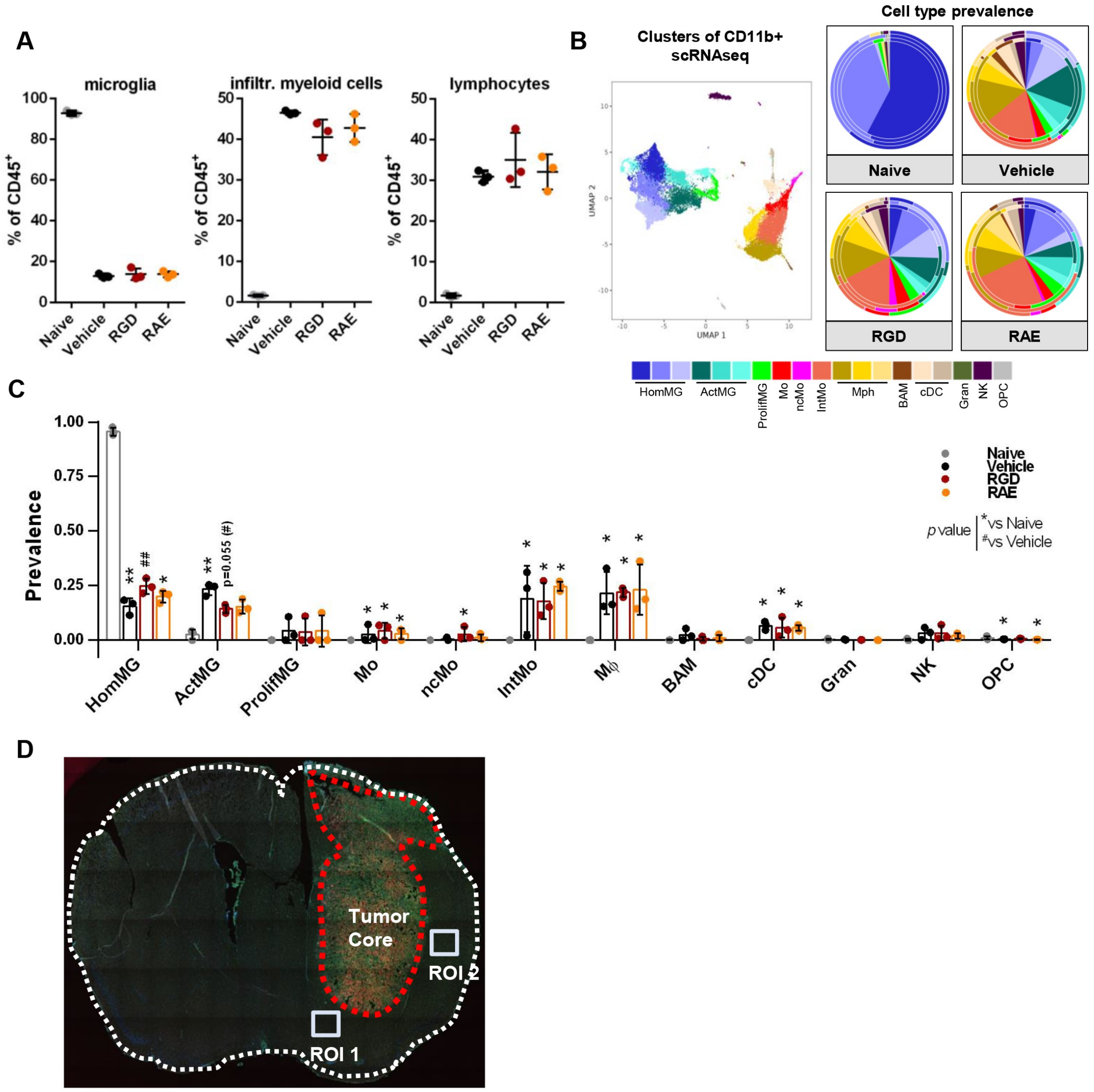


**Fig. S4. Profiles of myeloid cells in the brain of RGD-treated GL261 glioma-bearing mice.** (A) Percentages of microglia (CD11b$^{+}$CD45$^{low}$), infiltrating myeloid cells (CD11b$^{+}$CD45$^{high}$) and lymphocyte-enriched population (CD11b$^{-}$CD45$^{high}$) among immune cells isolated from vehicle ($H_2O$) and peptide-treated glioma-bearing mice at 21 DPI. Naive mice were used as a reference. (B, C) Cell type prevalence inferred by deconvolution of bulk RNA-seq data of sorted brain CD11b$^{+}$ cells. Deconvolution was based on transcriptomic signatures of different brain myeloid cell clusters (B, left panel) identified using CITE-seq in the same glioma model (*34*). Predicted cell type abundance is presented on pie-charts (B, right panel) with average values in the center and cell prevalence in each repetition indicated on the external rims. Bar plot (C) showing the prevalence of general cell types (values were pooled in case of multiple clusters per cell type) in each treatment condition. The signatures of activated microglia, intermediate monocytes, macrophages and classical dendritic cells are more abundant in $CD11b^+$ cells isolated from the tumor-bearing brains in comparison to higher prevalence of homeostatic microglia in naïve mice. Bars represent the mean ± SD. HomMG - homeostatic microglia, ActMG - activated microglia, ProlifMG - proliferating microglia, Mo - monocytes, ncMo - non-classical monocytes, IntMo - intermediate monocytes, Mφ - macrophages, BAM - border-associated macrophages, cDC - dendritic cells, Gran - granulocytes, NK - natural killer cells , OPC - oligodendrocyte progenitor cells. Significance was assessed with One–Way ANOVA and Uncorrected Fisher's LSD multiple comparison test; N=3; ***$p < 0.001$; **$p < 0.01$; *$p < 0.05$. (D) Regions Of Interest (ROI) for quantification of vessel density using lectin staining of the brain section.

### Supplementary Fig. 5

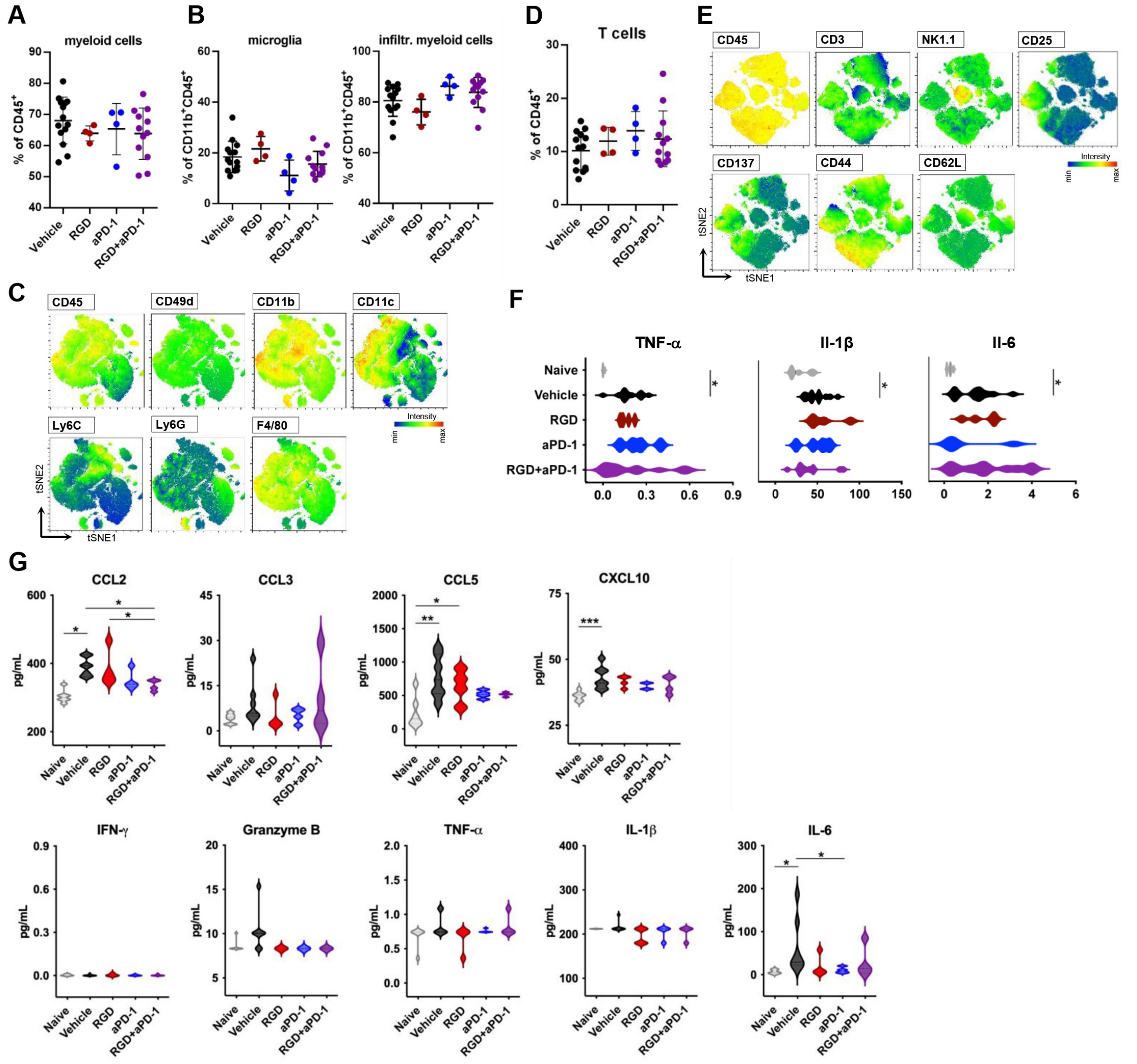


**Fig. S5. Characterization of the immune microenvironment of GL261 gliomas upon RGD or/and anti-PD-1 treatment.** Frequencies of: all CD11b+ myeloid cells (A), microglia ($CD11b^{+}CD45^{low}$) and infiltrating myeloid cells ($CD11b^{+}CD45^{high}$) (B), and $CD3^{+}$ T cells (D) in the glioma-bearing brain at 28 DPI upon a specific treatment. (C,E) Graphical visualization of the immune microenvironment using tSNE plots. Unsupervised tSNE clustering of $CD45^{+}CD11b^{+}$ isolated from brains of naive mice, Vehicle, RGD, anti-PD-1 (aPD1) and RGD+aPD-1 treated glioma-bearing mice. In each group, 40,000 cells were clustered, equal number from each replicate (obtained by down-sampling in FlowJo). tSNE plots depicting distribution of the markers discriminating myeloid subpopulations within $CD11b^{+}CD45^{+}$ cells (C) and lymphoid subpopulations within $CD11b^{-}CD45^{+}$(E) are presented. (F-G) The levels of pro/anti-inflammatory cytokines were determined in brain homogenates (F) and sera (G) of naive and treated mice at 28 DPI using a multiplexed Luminex bead-based assay. Violin plots show the levels of tested cytokines in pg/mg of total protein in homogenates and in pg/ml in sera. Significance was assessed with One–Way ANOVA and Uncorrected Fisher's LSD multiple comparison test; N=4-12; ***$p < 0.001$; **$p < 0.01$; *$p < 0.05$.

### Supplementary Fig. 6

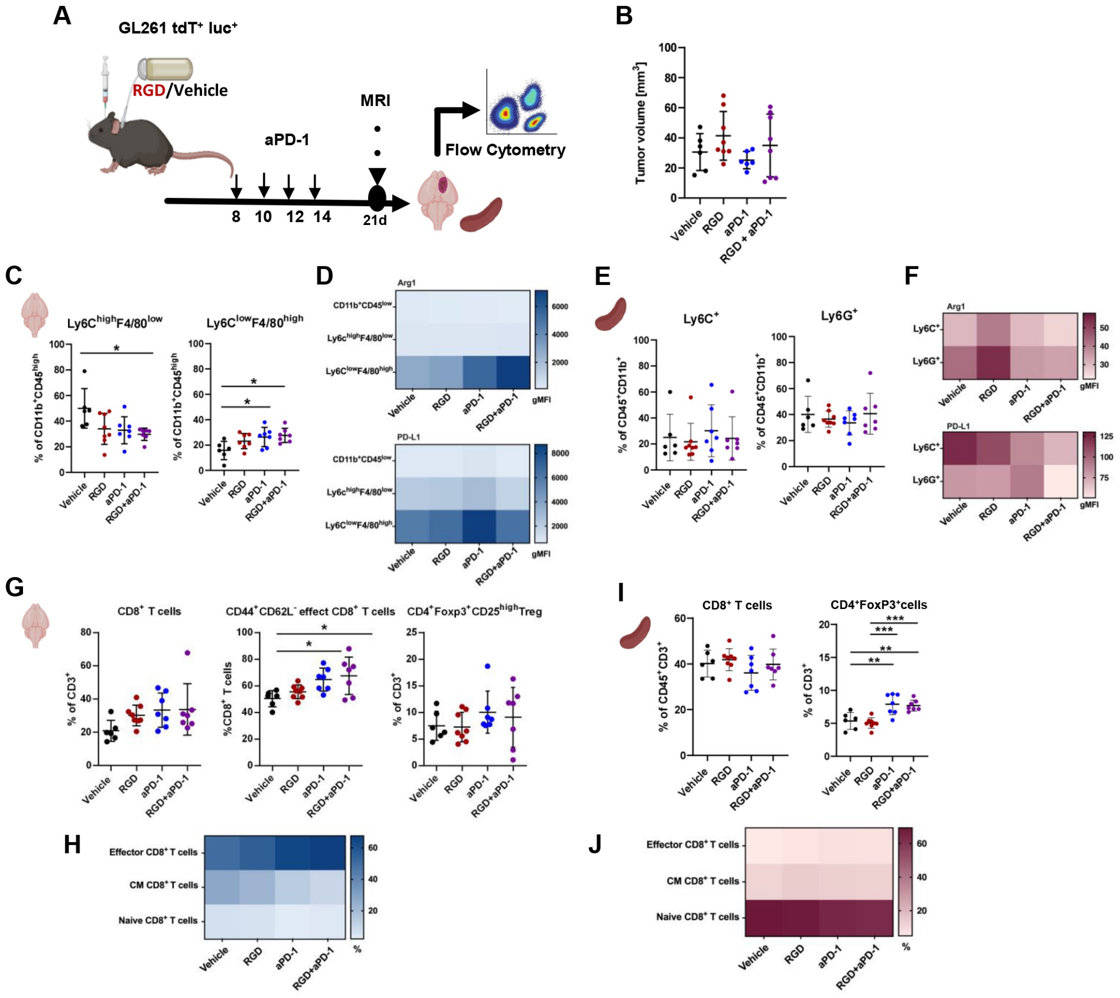


**Fig. S6. Immune cells phenotyping in glioma TME and spleens at 21 DPI in tumor-bearing mice upon administration of RGD+anti-PD-1**. (A) Experimental scheme depicting treatments and procedures. Mice were implanted with GL261 tdTomato+luc+ glioma cells and received water (Vehicle) or the RGD peptide via osmotic pumps and anti-PD-1 antibody (aPD-1, 10 mg/kg) or control IgG by i.p. injection at 8, 10, 12 and 14 DPI. (B) Quantification of tumor volume MRI at 21 DPI in mice; mean ±SD is presented; n=6-8 per group. At 21 DPI animals were perfused, tumor-bearing brains and spleens were removed and processed to isolate myeloid (C-F) and lymphoid cells (G-J) by FACS. Percentages of monocytes (Ly6C$^{high}$F4/80$^{low}$), and tumor associated macrophages (Ly6C$^{low}$F4/80$^{high}$) in the brains (C) and monocytes (Ly6C$^{+}$) and granulocytes (Ly6G$^{+}$) in the spleens (E) were evaluated. Heatmaps show levels (gMFI) of Arg1 and PD-L1 in these populations in the brains (D) and the spleens (F). (G,I) Percentages of CD8$^{+}$ T cells (CD8$^{+}$CD4$^{-}$), effector CD8$^{+}$ T cells (CD8$^{+}$CD44$^{+}$CD62L$^{-}$) and Treg cells (CD4$^{+}$Foxp3$^{+}$CD25$^{high}$) in the brains (G) and the spleens (I). Heatmaps shows percentages of $Ki67^{+}$ cells among effector ($CD44^{+}CD62L^{-}$), central memory (CM; $CD44^{+}CD62L^{+}$) and naïve ($CD44^{-}CD62L^{+}$) $CD8^{+}$ T cells in the brains (H) and the spleens (J). Statistical significance of differences between groups was assessed with One-Way ANOVA followed by Tukey's multiple comparison test. p-Values were considered as significant when $^{**}p < 0.01$; $^{*}p < 0.05$.

**Supplementary Fig. 7**

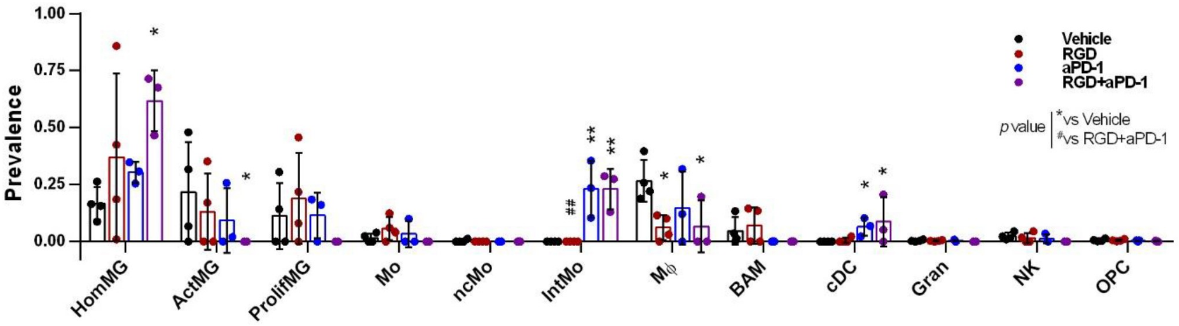


**Fig. S7. Myeloid cell profiles in the brains of RGD-, anti-PD-1 and RGD+anti-PD-1-treated GL261 glioma-bearing mice.** Cell type prevalence inferred from deconvolution of bulk RNA-seq data of CD11b$^{+}$ cells sorted from the brains of GL261 glioma-bearing mice at 28 DPI. Deconvolution was based on transcriptomic signatures of brain myeloid cell clusters (as in Fig. S4). Bar plot showing the prevalence of general cell types (values were pooled in case of multiple clusters per cell type) in each treatment condition. Bars represent the mean ± SD. For cell type description see Fig. S4. Significance was assessed with One–Way ANOVA and Uncorrected Fisher's LSD multiple comparison test; N=3; ***$p < 0.001$; **$p < 0.01$; *$p < 0.05$.

**Supplementary Fig. 8**

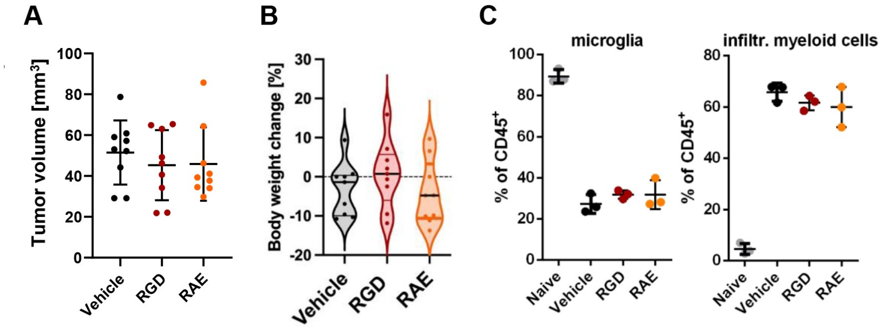


**Fig. S8. The response to RGD-treatment in U87MG-RFP+ glioma-bearing mice.** Athymic Nude-Foxn1nu mice were orthotopically implanted with U87MG-RFP$^{+}$ human glioma cells and received $H_2O$ (Vehicle), RGD or control peptide (RAE) via osmotic pumps. (A) Tumor volume was measured by MRI at 21 DPI. (B) Body mass of recipient mice (change from day 0 to21 DPI) in Vehicle and treated groups; n=9; the line is plotted at the mean. (C) Frequencies of microglia (CD11b$^{+}$CD45$^{low}$) and infiltrating myeloid cells (CD11b$^{+}$CD45$^{high}$) isolated from the brains of vehicle ($H_2O$) and peptide-treated U87MG-RFP+ glioma-bearing mice at 21 DPI. Naive mice were used as a reference.

**Supplementary Fig. 9**

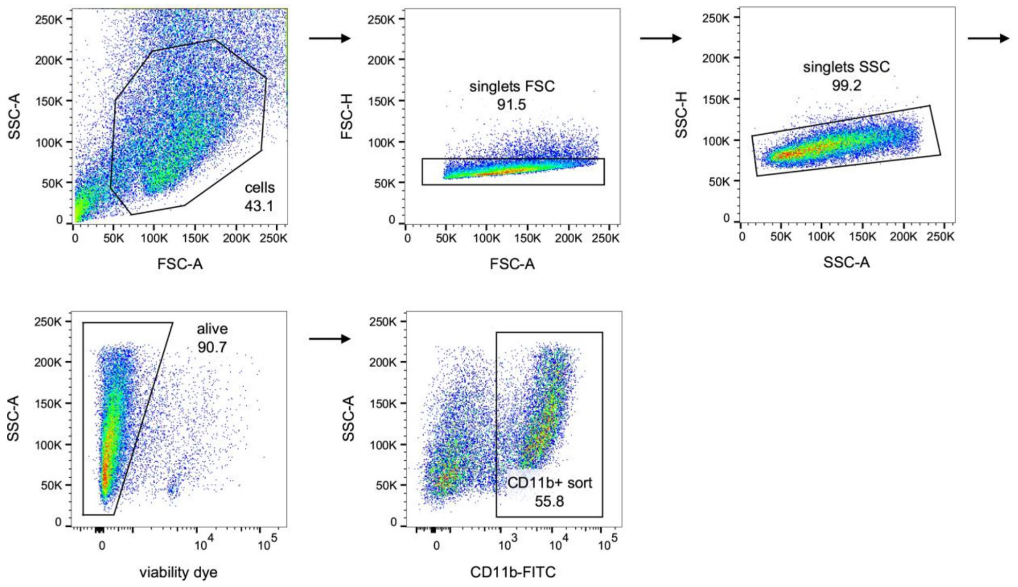


**Fig. S9. Gating strategy used for FACS-sorting.** Events corresponding to cells were gated on SSC-A vs FSC-A plots, then doublets were excluded. Events in singlets gate were further analyzed for the uptake of Fixable Viability Dye to exclude events corresponding to dead or damaged cells. CD11b$^+$ cells were selected for sorting on SSC-A vs CD11b plots.

### Supplementary Fig. 10

**A**

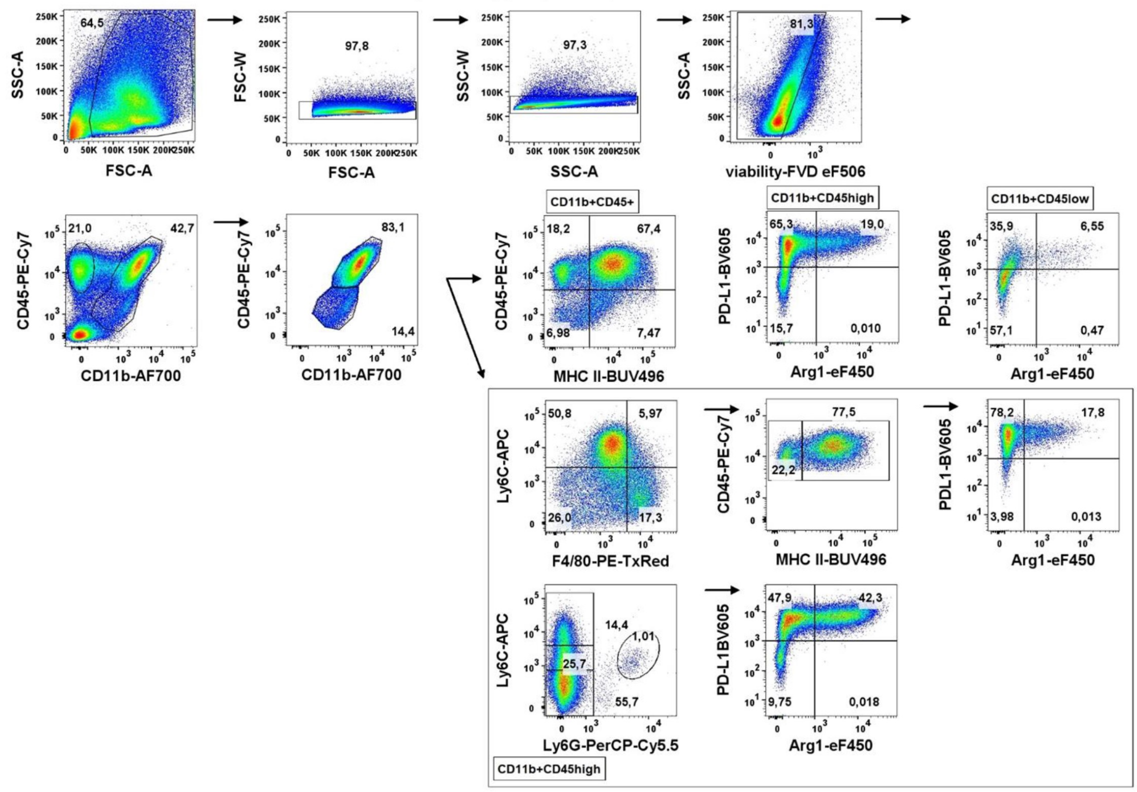


**B**

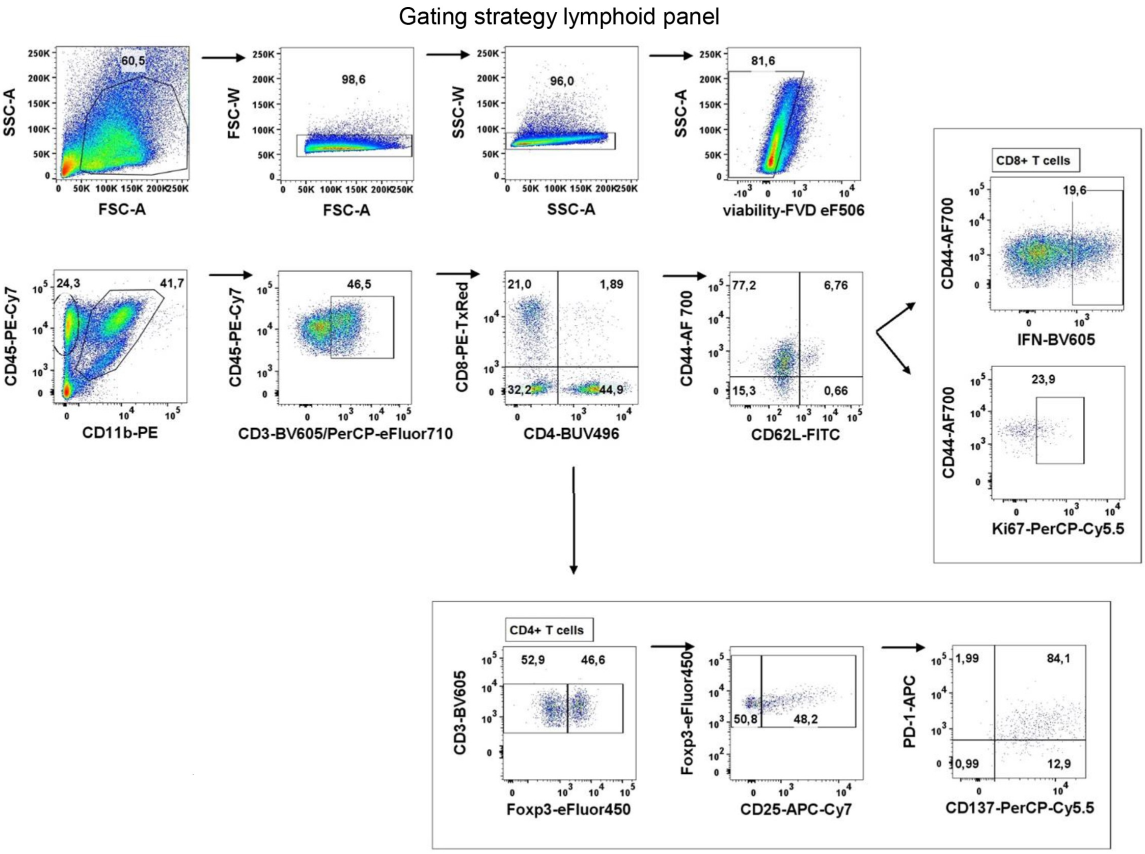

**Fig. S10. Gating strategy used for flow cytometry analysis.** (A) Representative gating strategy for analysis of (A) myeloid and (B) lymphoid cells. Events corresponding to cells were gated on SSC-A vs FSC-A plots, then doublets were excluded. Events in singlets gate were further analyzed for the uptake of Fixable Viability Dye to exclude events corresponding to dead or damaged cells. Quadrant gates were drawn on cell subpopulations based on differences in the surface expression of CD11b and CD45 antigens: microglia ($CD11b^{+}CD45^{low}$), blood-derived macrophages ($CD11b^{+}CD45^{high}$ in myeloid compartment and $CD3^{+}$, $CD4^{+}$ and $CD8^{+}$ in lymphoid compartment. Expression of markers associated with immune evasion (PD-L1, Arg-1) were verified using FMO controls.

### Tables

#### Table S1. Antibodies

| Antibodies | Company | Cat no. | Dilution |
|---|---|---|---|
| *Antibodies for flow cytometry* | | | |
| Arginase 1 Monoclonal Antibody (A1exF5) eFluor 450, | Life Technologies Corp. Carlsbad, CA, USA | 48-3697-82 | 1:100 |
| CD3e (145-2C11), Hamster Anti-Mouse, BV605 | BD | 563004 | 1:100 |
| CD8a Monoclonal Antibody (53-6.7) PE-eFluor 610 | Life Technologies Corp. Carlsbad, CA, USA | 61-0081-82 | 1:200 |
| CD4 Rat Anti-Mouse BUV496 | BD | 741051 | 1:200 |
| CD11b Rat Anti-Mouse (M1/70) FITC | BD | 553310 | 1:800 |
| CD11b Rat Anti-Mouse (M1/70) PE | BD | 553311 | 1:800 |
| CD11b Rat Anti-Mouse (M1/70) AF-700 | BD | 557960 | 1:800 |
| CD25 Monoclonal Antibody (PC61.5) APC-eFluor 780 | Life Technologies Corp. Carlsbad, CA, USA | 47-0251-82 | 1:50 |
| CD44 Monoclonal Antibody (IM7) Alexa Fluor 700 | Life Technologies Corp. Carlsbad,CA,USA | 56-0441-80 | 1:100 |
| CD45,Rat Anti-Mouse (30-F11) PE-Cy7 | BD | 561868 | 1:800 |
| CD49d Rat Anti-Mouse, FITC | Biolegend | 103606 | 1:400 |
| CD62L (L-Selectin) Monoclonal Antibody (MEL-14) FITC | Life Technologies Corp. Carlsbad, CA, USA | 11-0621-82 | 1:400 |
| CD137 Monoclonal Antibody (17B5) PerCP-eFluor 710 | Life Technologies Corp. Carlsbad, CA ,USA | 46-1371-82 | 1:100 |
| CD274 (PD-L1, B7-H1) Monoclonal Antibody (MIH5), Super Bright 600 | Life Technologies Corp. Carlsbad, CA, USA | 63-5982-82 | 1:100 |
| CD279 Hamster Anti-Mouse (J43) APC | BD | 562671 | 1:100 |
| F4/80 Monoclonal Antibody (BM8) PE-eFluor 610 | Life Technologies Corp. Carlsbad, CA, USA | 61-4801-82 | 1:100 |
| FOXP3 Monoclonal Antibody (FJK-16s) eFluor 450 | Life Technologies Corp. Carlsbad, CA,USA | 48-5773-82 | 1:100 |

| | | | |
|---|---|---|---|
| Ly6C Rat Anti-Mouse (AL-21) APC | BD | 560595 | 1:200 |
| Ly6G Rat Anti-Mouse (1A8) PerCP-Cy5.5 | BD | 560602 | 1:200 |
| MHC Class II (I-A/I-E) Rat Anti-Mouse (M5/114.15.2) BUV496 | BD | 750281 | 1:200 |
| Ki67 Mouse | Biolegend | 561284 | 1:200 |
| IFNγ, Rat Anti-Mouse (XMG1.2) BV605 | Biolegend | 505840 | 1:50 |
| *Antibodies for immunohistochemistry* | | | |
| anti-Arginase pAb | Novus Biologicals | NB100-59740 | 1:100 |
| anti-Iba1 | Wako | 019-19741 | 1:1000 |
| anti-rabbit Alexa Fluor 647 | Life Technologies Corp. Carlsbad, CA, USA | A31573 | 1:1000 |
| anti-goat Alexa Fluor 488 | Life Technologies Corp. Carlsbad, CA, USA | A11055 | 1:1000 |
